# Cuprizone drives divergent neuropathological changes in different mouse models of Alzheimer’s disease

**DOI:** 10.1101/2023.07.24.547147

**Authors:** Gerald Wai-Yeung Cheng, Iris Wai-Ting Ma, Jianpan Huang, Sunny Hoi-Sang Yeung, Paolo Ho, Zilin Chen, Henry Ka Fung Mak, Karl Herrup, Kannie Wai Yan Chan, Kai-Hei Tse

**Author notes:** Send correspondence to: Kai-Hei Tse PhD, Department of Health Technology and Informatics, Lee Shau Kee Building, The Hong Kong Polytechnic University, Hung Hom, Kowloon, Hong Kong SAR. Contributed equally (Gerald Wai-Yeung Cheng, Iris Wai-Ting Ma, Jianpan Huang).

## Abstract

Myelin degradation is a normal feature of brain aging that accelerates in Alzheimer’s disease (AD). To date, however, the underlying biological basis of this correlation remains elusive. The amyloid cascade hypothesis predicts that demyelination is caused by increased levels of the β-amyloid (Aβ) peptide. Here we report on work supporting the alternative hypothesis that early demyelination is upstream of amyloid. We challenged two different mouse models of AD (R1.40 and APP/PS1) using cuprizone-induced demyelination and tracked the responses with both neuroimaging and neuropathology. In oppose to amyloid cascade hypothesis, R1.40 mice, carrying only a single human mutant APP (Swedish; APP_SWE_) transgene, showed a more abnormal changes of magnetization transfer ratio and diffusivity than in APP/PS1 mice, which carry both APP_SWE_ and a second PSEN1 transgene (delta exon 9; PSEN1_dE9_). Although cuprizone targets oligodendrocytes (OL), magnetic resonance spectroscopy and targeted RNA-seq data in R1.40 mice suggested a possible metabolic alternation in axons. In support of alternative hypotheses, cuprizone induced significant intraneuronal amyloid deposition in young APP/PS1, but not in R1.40 mice, and it suggested the presence of PSEN deficiencies, may accelerate Aβ deposition upon demyelination. In APP/PS1, mature OL is highly vulnerable to cuprizone with significant DNA double strand breaks (53BP1^+^) formation. Despite these major changes in myelin, OLs, and Aβ immunoreactivity, no cognitive impairment or hippocampal pathology was detected in APP/PS1 mice after cuprizone treatment. Together, our data supports the hypothesis that myelin loss can be the cause, but not the consequence, of AD pathology.

**SIGNIFICANCE STATEMENT:** The causal relationship between early myelin loss and the progression of Alzheimer’s disease remains unclear. Using two different AD mouse models, R1.40 and APP/PS1, our study supports the hypothesis that myelin abnormalities are upstream of amyloid production and deposition. We find that acute demyelination initiates intraneuronal amyloid deposition in the frontal cortex. Further, the loss of oligodendrocytes, coupled with the accelerated intraneuronal amyloid deposition, interferes with myelin tract diffusivity at a stage before any hippocampus pathology or cognitive impairments occur. We propose that myelin loss could be the cause, not the consequence, of amyloid pathology during the early stages of Alzheimer’s disease.

## INTRODUCTION

Neuroimaging studies have identified myelin degradation as one of the earliest events in the progression of sporadic Alzheimer’s disease (AD)(1). Studies using a range of magnetic resonance imaging (MRI) techniques including diffusion kurtosis/tensor (DKI/DTI) (2), magnetization transfer (MT) (3) and magnetic resonance spectroscopy (MRS) (4) have shown that myelinated tracts progressively degrade at during normal aging. The timing of such age-related myelin loss coincides with the decline of peak cognitive performance (5) and is strongly correlated with an increase of β-amyloid and phosphorylated tau in the cerebrospinal fluid of an AD patient cohort (6). The oligodendrocyte (OL) is the only cell type that myelinates axons in the central nervous system and both its survival and differentiation are vulnerable to amyloid peptides (7, 8). We and others reported that the loss of OL in AD brain can be recapitulated in distinct transgenic mouse models carrying single (*APP* (9)), double (*APP*, *PSEN1* (10)) and triple (*APP*, *PSEN1*, *MAPT* (11–13)) genetic mutations associated with the amyloid cascade hypothesis. Based on these findings, the simplest hypothesis would be that amyloidosis causes myelin degradation in the aging brain, and that the more severe amyloid pathology will be associated with the more severe myelin pathology.

As compelling as amyloid cascade hypothesis might be, there is at least one alternative amyloid-independent hypothesis exists to explain the myelin loss found in the aging brain. The earliest reduction of intracortical myelination is found in the frontal lobe, and such loss is closely correlated with cognitive impairment (14, 15). The natural age-related loss of myelin begins in the fifth decade of life in healthy population (16). This timing is well before the formation of significant amyloid aggregation in the aging brain or changes in cerebrospinal fluid or serum levels of Aβ (17). Indeed, once the symptoms of dementia become apparent, the accelerated reduction of OL is found regardless of either the levels of amyloid plaque (9), or the presence or absence of tau deposits (i.e. Braak Stage) (18).

A nuanced perspective of the situation arises from considering that myelinating OLs are the primary target of ischemia, inflammation, amyloidosis, oxidative stress, and DNA damage (19–21). As all these toxic factors gradually accumulate in the aging brain, the age- and disease-related myelin loss is likely the consequence of the summation of these diverse forces acting on OL lineage. In turn, this begins a vicious cycle as OL loss leads to multiple neuronal deficiencies including saltatory conduction disruptions and axonal transport abnormalities (22–25). The latter would have important implication for AD because the amyloid precursor protein (APP) is continuously moved by axonal transport (26, 27) to serve many physiological functions (28). Compromised axonal transport and the subsequent axonal injury that occurs upon demyelination (29–31), may combine to alter APP dynamics. This alternative hypothesis paints a more realistic picture where the insults of age and damage, secondary to myelin and OL loss, causes APP accumulation and accelerate their cleavage into β-amyloid (32, 33). In this scenario myelin degradation presents as a cause, rather than a consequence, of amyloid pathology (17, 34).

Here, we tested the above hypothesis in two AD mouse models, R1.40 and APP/PS. In APP/PS1, carrying with APP_SWE_ and presenilin 1 transgenes (PSEN1_dE9_), diffuse amyloid plaque formation is detectable by 6 months old (35–37). But in R1.40, carrying APP_SWE_ transgene alone, diffuse plaque does not form until 14 months old (38). Such amyloidosis difference was shown to translate into distinct onset age of cognitive impairments (39). After challenging young animals with cuprizone before any plaque is formed, we showed significant differences between the two models in the myelin tract integrity (DTI), myelin content (MTR) and metabolic shifts (MRS). These differences were independent of their severity of amyloidosis. Importantly, demyelination induced intraneuronal amyloid deposition formation in APP/PS1 – an early neuropathology in human AD brain (40, 41). These findings offer a mechanism by which OL deficiencies trigger amyloid pathology rather than the reverse.

## MATERIALS AND METHODS

### Animal subjects

All animal experiments were approved by the Animal Subjects Ethics Sub-Committee at The Hong Kong Polytechnic University (PolyU) and all experiments involving animals were performed under a valid license from the Department of Health of Hong Kong government in compliance with the Ordinance and The Code of Practice for Care and Use of Animals for Experimental Purposes of Hong Kong. All animals were housed in a temperature- and humidity-controlled environment with individually ventilated cage in a dedicated room on a 12-hour light/dark cycle with food and water *ad libitum*. All mice were kept and cared for at Centralized Animal Facilities (CAF) at PolyU.

Transgenic AD mouse models, R1.40 and APP/PS1, were obtained from The Jackson Laboratory (Bar Harbor, Maine, US). R1.40 (B6.129-Tg(APPSw)40Btla/Mmjax; MMRRC Strain #034831-JAX) was created by the insertion of the entire human amyloid precursor protein APP with the Swedish mutation (K670N/M671L; APP_SWE_) into the mouse genome on a yeast artificial chromosome transgene (35). These animals express all isoforms of APP_SWE_ at mRNA level at about 3-folds higher than that of endogenous mouse *App.* Significant diffuse plaques first appear in frontal cortex at 14 months old on the C57BL/6 background (42). APP/PS1 (B6;C3-Tg(APPswe,PSEN1dE9)85Dbo/Mmjax; MMRRC Strain #034829-JAX) is a double transgenic mouse model with a chimeric mouse/human APP gene with the Swedish mutation plus a human presenilin (PSEN1) with exon-9-deleted mutation (PSEN1_dE9_). This double-transgene was inserted at a single locus on chromosome 9 and under control of the mouse *Prnp* promoter and APP/PS1 animals exhibit diffuse plaque deposits as early as 6 months of age (43). Hemizygote animals were employed for experiments in both strains, and the non-carriers were employed as wildtype (WT) controls.

### Cuprizone-mediated demyelination

Male mice between 8 – 12 weeks old were subjected to cuprizone-mediated demyelination before experiments described below. The decision to use only male animals was because there is well-known gender-specific effect of cuprizone on female rodents (44, 45). Briefly, animals were fed with a cuprizone containing diet (CPZ; 0.2% w/w, TD.00588, irradiated, Envigo, Indianapolis, IN, US) or with a control diet (TD.140803, irradiated, Envigo, Indianapolis, IN, US) for four weeks *ad libitum*. The consumption of the diet was monitored on alternate days and no difference was observed among genotypes. No adverse effects on motor function nor mortality were observed throughout the experiment.

### Behavioral tests

All animals were randomized after cuprizone treatment and subjected to a Y-maze based behavioral test to assess spatial memory in CAF (46, 47). A custom-made opaque acrylic Y-Maze with three arms measuring 35(L) × 9(H) × 7(W) cm set at 120° angles apart was placed at the center of a room with dim lighting. All subjects were acclimatized for at least 1 hour before testing. The ANY-maze system with camera, computer, and software (version 7.2, Stoelting Co. IL, USA) was used for animal tracking. To assess spatial working memory, the spontaneous alteration was measured. The subject was placed at the center of the arms and allowed to freely roam the maze over the course of 8 minutes. Due to the new environment. The subject is expected to exhibit a tendency to enter a new arm (alternations) than a to enter a more recently visited arm (non-alternations). Spontaneous alternation is defined as the entries of all three arms on consecutive occasions and calculated as

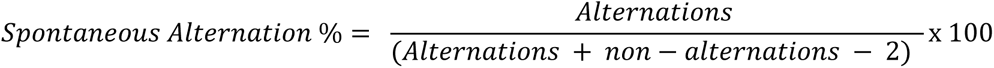

To assess spatial reference memory, the subjects were subjected to a training trial where one arm of the maze was blocked. The animals were allowed 4 minutes to freely explore the remaining two arms before being returned to their home cage. This blocked arm is designated as the correct arm for the trial. After inter-trial interval of one hour, the animal was placed back in the Y-Maze with all arms unblocked for 4 minutes. The number of entries and the time spent to the correct arm were assessed as the measure of spatial reference memory. Other behavioral parameter measurements were made by ANY-maze software included time spent in each arm, time spent in the correct arm, time spent in the incorrect arm, total entries, total distance travelled, average speed of the mouse, and average unmoving time. A significant reduction of spontaneous alternation and/or correct arm entries will be considered as indicator of impaired hippocampus function.

### Magnetic Resonance Imaging

All MRI experiments were conducted at the City University of Hong Kong (CityU) with a horizontal bore 3T Bruker BioSpec system (Bruker, Ettlingen, Germany) equipped with an 82 mm quadrature volume resonator as a transmitter and a single surface coil as a receiver.

Briefly, mice were initially anesthetized using a chamber with 2% isoflurane in oxygen generated by an oxygen concentrator (DeVilbiss Healthcare, Somerset, USA). After successful anesthesia induction, the animal was placed on a mouse cradle in the scanner with a gas inhalation mask to maintain anesthesia with 1–1.5% isoflurane. Mouse body temperature was maintained at 37°C using a warming pad connecting to a water heating system (Thermo Scientific, Waltham, USA). Respiration was continuously monitored using an MRI-compatible monitor system (SA Instruments, Stony Brook, USA). The image slices were positioned based on a collected sagittal image of the mouse brain with the position of the central coronal image slice set to -1.5 mm with respect to the anterior commissure (AC). For each mouse, T1/T2 weighted (T1w/T2w) imaging, diffusion tensor imaging (DTI) and magnetization transfer ratio (MTR) imaging and magnetic resonance spectroscopy (MRS) were acquired with a total scan time of about 37 minutes. For imaging acquisitions, eleven image slices with a matrix size of 128 ×128, a field of view (FOV) of 20 × 20 mm and a thickness of 1 mm were acquired. Other acquisition parameters for each sequence were specified as followings:

#### T1/T2 weighted imaging

Both T1w and T2w images were acquired using a rapid acquisition with refocused echoes (RARE) sequence. T1w parameters were set as: repetition time (TR) = 500 ms, echo time (TE) = 14 ms, number of average (NA) = 8, RARE factor = 4 (linear encoding), scan time = 2 min 8 sec. T2w parameters were set as: TR = 3000 ms, TE = 54 ms, NA = 8, RARE factor = 16, scan time = 2 min 24 sec.

#### DTI imaging

Diffusion tensor images were acquired using a spin echo-echo planar imaging (SE-EPI) sequence. Imaging parameters were set to: TR = 3000 ms, TE = 36 ms, NA = 4, diffusion encoding δ = 3 ms and diffusion separation Δ = 16 ms), 30 diffusion directions with b = 1000 s/mm^2^ and 5 unweighted b = 0 s/mm^2^ images, 2 experiments per direction, resulting in a scan time of 13 min. The diffusion tensor eigenvalues (λ1, λ2, λ3) computed by the ParaVision 6 were used to calculate parameter maps on a pixel-by-pixel basis (1) for the axial diffusivity (AD) with AD = λ1, the radial diffusivity (RD) with RD = (λ2 + λ3) / 2, the mean diffusivity (MD) with MD = (λ1 + λ2 + λ3) / 3 and the fractional anisotropy (FA) (2) with:

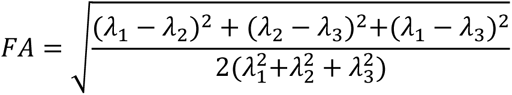

#### Magnetization Transfer Ratio

Magnetization transfer (MT) images were acquired by a MT sequence composed of an off-resonance radiofrequency (RF) irradiation as a saturation module and a RARE sequence as a readout module (48, 49). To compute the MT Ratio (MTR) maps, which are semi-quantitative measurements of macromolecules via MT effect, images were acquired at saturation offsets (ω) of ± 50, ± 25, ± 12.5, ± 5, ± 2.5 and 0 ppm using a continuous-wave (CW) RF saturation pulse with a saturation power (B_1_) of 2 μT and a saturation time (t_sat_) of 3 s. Imaging parameters were set to: TR = 40477 ms, TE = 7 ms, NA = 1, RARE factor = 96 (linear encoding).scan time = 9 min 26 sec. Images were processed using custom-written MATLAB (MathWorks, USA) scripts with MTR = (S_0_ – S_sat_(ω)/S_0_, where S_0_ and S_sat_(ω) are the signal amplitude measured without and with a saturation pulse at frequency ω, respectively.

#### Magnetic Resonance Spectroscopy

A voxel of interest (VOI) for MRS acquisition was localized in cortex region of each mouse with reference to the acquired T2 weighted image. Localized shimming up to second order was performed using voxel-based field mapping before each MRS acquisition. MRS data were acquired using the stimulated echo acquisition mode (STEAM) sequence with outer volume suppression. The acquisition parameters were voxel size = 3 × 3 × 3 mm^3^, FID = 2048 points, averages = 256, TE = 3ms, mixing time (TM) = 10 ms, TR = 2300 ms, scan time = 9 min 49 s. The location of the volume encompasses the cortex, corpus callosum, hippocampus, and thalamus between Bregma -1.3 to -3.2 (mm). Water suppression was achieved using the variable pulse power and optimized relaxation (VAPOR) method. MRS data processing and analysis were performed in MATLAB (MathWorks, USA). MRS were aligned by setting the peak of N-acetyl aspartate (NAA) at 2.02 ppm (50). For quantification of metabolites, the resonance integral of creatine (Cr, 3.11-2.92 parts per million (ppm)) against that of myo-inositol (mI, 3.62-3.47 ppm), choline (Cho, 3.31-3.11 ppm), creatine Glx (glutamine + glutamate, 2.52-2.24 ppm), NAA (N-acetyl aspartate, 2.24-1.88 ppm), methylene-group containing lipid/macromolecule (CH_2_-Lipid, 1.55-1.06 ppm) and methyl-group containing lipid/macromolecule (CH_3_-Lipid, 1.06-0.73 ppm) was calculated (51), and ratio was presented.

#### Radio-pathological investigation

In all MRI experiment, except for MRS, the whole brain was scanned. The regions of interests (ROI) including cortex, anterior cingulate cortex, hippocampus, corpus callosum, cingulum and external capsule spanning across Bregma -3.4 to 2.7 were traced and annotated offline in MATLAB for data extraction. At the end point of each experiment, the animals were deep anesthetized by Avertin (1.25% tribromoethanol, 375 mg/kg, intraperitoneal) before transcardial perfusion with phosphate buffer saline (PBS) using a peristaltic pump as described before (52). The animal was then dissected, and its brain was isolated. The left hemisphere was snap frozen for protein and gene expression analysis, while the right hemisphere was fixed by immersion in paraformaldehyde (4%) for 24 hours at 4 °C for histological analysis. The fixed tissue was cryopreserved and dehydrated in sucrose (30% w/v in PBS, 4°C) for 48 hours. To isolate atlas-referenced brain sections to match MRI-findings for analysis, the processed brain hemisphere was sliced through a metal cast brain matrix for adult mouse (coronal, 1 mm interval, #15003, Ted Pella Inc, CA, USA). These brain slices between Bregma -3.4 to 2.7 were mapped radiologically and histologically against using Allen Mouse Brain Atlas (53). The Bregma-specific brain slices were then embedded for cryosectioning at 10 μm in coronal planes. All coronal sections were mounted on glass slides and kept at -80 ^°^C until analysis.

### Targeted RNA-seq

#### RNA extraction and targeted RNA sequencing

Whole cerebral cortices were isolated from mouse brain tissue samples for RNA extraction. The hippocampus and residue WM tissue were removed by microdissection. Tissue homogenization was performed with the Precellys Evolution Homogenizer (Bertin Technologies, France, P000062-PEVO0-A) at 5500 rpm for 3 cycles of 10s with a 10s pause, and tissues were lysed in TRIzol (Thermo Fisher, United States, 15596026) simultaneously. RNA was isolated from homogenate with RNeasy Mini spin columns (Qiagen, Germany, 74106). Extracted RNA was quantified and qualified with Agilent RNA 6000 Nano Kit using Agilent Bioanalyzer. A total of 400 ng an RNA with RNA integrity number (RIN) greater than 8 from each sample was processed with QIAseq Targeted RNA Mouse Cancer Transcriptome Panel (Qiagen, Germany, RMM-003Z). This panel consists of 416 genes selected for the association with angiogenesis, apoptosis, cell cycle, cell senescence, DNA repair and damage, epithelial-to-mesenchymal transition, hypoxia, metabolism, telomeres and telomerase, plus 14 reference normalization genes. The library preparation was performed according to manufacturer’s instruction. cDNA was synthesized from each RNA sample and each cDNA molecule was then barcoded (BC) with an unique molecular tag (MT) primer, gene-specific sequences (GS) and common Universal primer sequence (RS2). Unique MT attachment was used to identify the original number of cDNA gene reads for normalization on gene expression. Barcoded cDNA was then purified with QIAseq beads before PCR amplification with gene-specific primers. Target amplicons of each sample were amplified universally and indexed with QIAseq Targeted RNA 12 index I. After a final cleanup with QIAseq beads, libraries of each sample were analyzed with Agilent High Sensitivity DNA Kit. A total of six libraries of samples were pooled at 12 pM with 1% PhiX spike-in and paired-end sequenced with Illumina MiSeq system using Illumina MiSeq Reagent Kit v3 (150-cycle) (Illumina, United States, MS-102-3001). An average of 5.60 Gb data was obtained from 1849.27 K/mm2 mean cluster density in each sequencing run on MiSeq. On average, 91.15% of the clusters (34.18 million counts) passed the quality control filter at Q30.

#### Analysis of targeted gene expression

FastQ files of each library with an average of 4.6 million read depth were uploaded to Qiagen’s GeneGlobe online portal for gene expression quantification. According to the internal pipeline of QIAseq RNA amplicon read processing, universal PCR adapter oligos and molecular tag (MT) sequences were trimmed from reads before alignment to GRCm38 reference genome using the STAR RNA read mapper. Reads with their alignment length shorter than 60 base pairs or demonstrating poor alignment at primer start sites were dropped. Aligned reads were then clustered according to MT regions for read count on each amplicon. Counts of MT read clusters on each targeted region were normalized with geNorm. Averaged unique molecular index counts of ten genes with best geNorm-calculated stability factors less than 1.5 were selected from the entire gene set as the normalization factor. Correlation among samples was determined by with Principal Component Analysis (PCA) and a Pearson correlation heatmap, which were both generated by ggpubr (v 0.6.0) and ggheatmap (v 2.2). P-values of false discovery rate (FDR) from grouped comparisons were calculated with Benjamini-Hochberg method and resulted in volcano plots with the log-fold-change generated by EnhancedVolcano (v 1.13.2) library. Standardization of z-score was performed on 55 out of 416 differentially expressed genes (DEG) A differential expression heatmap was plotted using z-score of gene expressions and the GO mapping of each significant DEG generated by ComplexHeatmap (v 2.14) library. The upregulated and downregulated genes with a P value < 0.05 were identified and then examined by PANTHER-based gene ontology (GO) enrichment analysis (Gene Ontology Consortium) (54–56), against the Qiaseq panel gene list.

### Immunohistochemistry

Histological analysis was performed with immunohistochemistry using the fully automated stainer BOND RX ® system (Leica Biosystems GmbH, Germany). Using BOND Polymer Refine Detection Kit (Leica Biosystems, IL, USA) as described (52). The sections were washed, rehydrated, and incubated with antibodies against OL markers proteins including Olig2 (MABN50, 1:500 Millipore, CA, USA, heat induced epitope retrieval-(HIER) with ER1 solution for 20 min), Nkx 2.2 (Abcam, ab191077, 1:1000 Abcam, Cambridge, HIER with ER2 solution for 20 min), or APC (Clone CC1, MABC200, 1:500 Millipore, CA, USA HIER with ER1 solution for 20 min), PLP1 (Clone E9V1N, CST 28702S, 1:500, HIER with ER1 solution for 20 min), MOG (Clone E5K6T, CST 96457S, 1:1000, Cell Signaling Technology, MA, USA, HIER with ER2 solution for 20 min). The amyloid marker, anti-β-Amyloid 1-16 (Clone 6E10, 803014, 1:5000, BioLegend, CA, USA, HIER with ER1 solution for 20 min) was used to detect amyloid depositions; The DNA damage marker, 53BP1 (ab36823, 1:5000 Abcam, Cambridge, UK, HIER with ER2 solution for 20 min) was used to detect genomic integrity. βIII-Tubulin (βIIItub)(1:10000, Clone TUJ1, 801201 BioLegend, CA, USA) was used as an axonal and neuronal marker. Primary antibody incubations were followed by signal detection using BOND Polymer Refine Detection Kit (Leica Biosystems, IL, USA) with Mayer’s hematoxylin nuclear counter stain as per manufacturer’s instructions. All sections were then coverslipped for microscopic examination and digital pathology.

### Whole slide image acquisition

Whole slide images (WSI) of each batch of stained brain tissues at Bregma 0.8 and -1.3 were acquired at 40x magnifications by either Leica Aperio CS2 digital pathology slide scanner (resolution at 0.252 µm/ pixel, Leica Biosystems, Hong Kong) or NanoZoomer S210 digital slide scanner (resolution at 0.221 µm/ pixel, Hamamatsu Photonics K.K., Japan). No difference was found between the WSI acquired from the two system, as reported earlier (57).

### Whole slide image quantifications

WSIs were analyzed by an open-source software platform for quantitative pathology analysis, QuPath v0.3.2 (58). QuPath has been employed for experimental neuropathology and clinical cancer diagnosis (59, 60). The source code, documentation, and full software of QuPath are freely accessible at https://qupath.github.io. Briefly, WSIs of the same markers were loaded into QuPath as a project and imported as a H-DAB image for automatic colour deconvolution into blue (hematoxylin), brown (DAB) and residual colour. ROIs identified in MRI investigation were further refined and segmented on each WSI in QuPath according to Allen Mouse Brain Atlas. Prior to any immunohistochemistry analysis, the cellularity of the image was first evaluated by built-in Cell Detection tool, which automatically segments the cell (nucleus) from surroundings. The difference between neuronal and non-neuronal cells was first assessed by the difference in cell size and circularity in the cortex and the white matter (genu), where the majority of constituent cell types are oligodendrocytes and astrocytes. Cellularity information acquired from at least three independent section was employed to optimize the parameters (pixel size, background radius, median filter radius, sigma, minimum & maximum area, and haematoxylin intensity threshold) of cell detection in immunohistochemistry. The optimized parameters were then evaluated for the segregation of neuron and non-neuronal cells in the gray matter by on a trial-and-error basis. The final cell segmentation parameters were correlated with manual counting done by the experimenter. The results were then used with image of other structures to distinguish neuronal and non-neuronal cells. The perimeters of requested pixel size, background radius, median filter radius, sigma, and the minimum & maximum area were kept constant among all specimen and the threshold values for hematoxylin and DAB were optimized by the experimenter on a batch-by-batch basis. Each batch of WSIs were analyzed by QuPath with a script without interference from the experimenter. The cell detection and annotation data were extracted from QuPath for downstream calculations, sorting and analysis in spreadsheets, and the samples were ublined only after the enterie analysis was completed. For axonal changes, three blinded investigators ranked randomized images of the cortex with βIII-tubulin immunohistochemistry on a score of 1 to 5, with 1 being the lowest and 5 being the highest axonal density.

### Immunoblotting

The biochemical changes in the whole cortex were examined by immunoblotting. Briefly, the entire frozen neocortex of the left hemisphere was grinded by Dounce homogenizer in RIPA buffer (radioimmunoprecipitation assay buffer, 20-188, Millipore, CA, USA) in the presence of protease inhibitors and phosphatase (Roche, Hong Kong). Protein samples were then quantified for normalization (Bio-Rad Protein Assay, 5000001, Bio-Rad, CA, USA). A total of 30 μg proteins were loaded per lane and electrophoresed on a 8 to 12% Sodium dodecyl sulfate-polyacrylamide gel. The resolved proteins were then transferred to polyvinylidene difluoride (PVDF) membranes using semi-dry method on Trans-Blot Turbo Transfer System (1704150, Bio-Rad, CA, USA). Any potential non-specific bindings were blocked by 5% non-fat milk for half hour before incubation with primary antibody. Primary antibodies specifically against β-Amyloid, 1-16 Antibody (1:000, Clone 6E10, N-terminal, a.a. 1-16, human specific only, 803014 BioLegend, CA, USA); β-Amyloid, 17-24 Antibody (1:1000, Clone 4G8, N-terminal, a.a. 17-24, human & mouse specific, 800708 BioLegend, CA, USA); Amyloid Precursor Protein (1:1000, Clone Y188, C-terminal, a.a. 750-770, human & mouse specific, Abcam, Cambridge, UK); βIII-Tubulin (βIIItub)(1:2500, Clone TUJ1, 801201 BioLegend, CA, USA); MOG (1:1000, MAB5680, Millipore, CA, USA and GAPDH (Loading control, ab8245, 1:10000, Abcam, Cambridge, UK). PLP (1:1000, PA3-150, Thermo Fisher Scientific, MA, USA) where then applied. The probed membranes were incubated with horseradish peroxidase-linked secondary immunoglobulins (Cell Signaling Technology, MA, USA). After washing with tris-buffer saline-Tween 20, the signals were visualized by chemiluminescent substrates (SuperSignal™ West Pico or Extended Dura, Thermo Fisher Scientific, MA, USA) and image captured by continuous exposure for 7 minutes with a Bio-Rad ChemiDoc MP Imaging System (12003154, Bio-Rad, CA, USA). The protein expression was quantified by determining individual band density using ImageJ (FIJI, v1.53t, National Institute of Health, MD, USA) with GAPDH as the loading control.

### Gene expression analysis

The gene expression analysis of the fixed brain sections was performed using real-time polymerase chain reaction. The total RNA from the fixed frozen brain tissues was isolated by PureLink FFPE RNA Isolation Kit (K156002, Thermo Fisher Scientific, MA, USA). All contaminating genomic DNA was removed by DNase I (1 U/μL, Thermo Fisher Scientific, MA, USA). Reverse transcription of the resulting RNA was performed using High-capacity RNA-to-cDNA kit (Applied Biosystem, Thermo Fisher Scientific, MA, USA) and the expression of specific gene was analysed using TB Green® Premix Ex Taq™ II (TaKaRa Bio Inc., Japan) in a Roche Light Cycler 480 with predesigned and validated primers from PrimerBank of Massachusetts General Hospital (61). All gene expression levels were calculated relative to wildtype control using housekeeping genes (GAPDH) based on the 2^-ΔΔCt^ method (62).

### Statistics

In power analysis, the sample size in the experiment design was estimated *a priori* using G*Power package (Version 3.1.9.7) (63). At least three independent biological replicates were included in each experiment and the *n* number is denoted in the graphs as the number of independent animal subjects employed. If necessary, a Grubbs’ test was used to identify, if any, outliers are presence. In the statistical analysis, comparisons between two groups were assessed using unpaired t-tests were performed. For comparisons of more than two groups, one-way analysis of variance (ANOVA) followed by post-hoc multiple comparison tests were performed. Two-way ANOVA followed by post-hoc multiple comparison tests was performed for two-independent variable analysis across genotype or treatment. All statistical tests are two-tailed, and the significance level was set at p < 0.05. The statistical test and post-hoc test in the multiple comparisons are stated in each figure legend. All statistical analysis were conducted using GraphPad Prism software version 9.50 (GraphPad Software Inc. MA, USA). The data points in the graphs are plotted as the mean ± standard errors.

## RESULTS

### Differences in MTR-detected myelin content and demyelination response between R1.40 and APP/PS1

Demyelination by feeding mice a cuprizone (CPZ) diet is a standard rodent model for multiple sclerosis (64–66). We used a four-week protocol to induce acute demyelination in wildtype (WT), R1.40 and APP/PS1 as illustrated in Fig. 1A. In all mouse strains, demyelination in both gray matter (GM) and white matter (WM) was achieved after four weeks, as illustrated by the immunohistochemistry of PLP (Fig. 1B), and in agreement with earlier report (65, 67). Yet, in T2w neuroimaging, only minimal differences were observed across mouse strains (Fig. 1C, top row).

**Figure 1.**
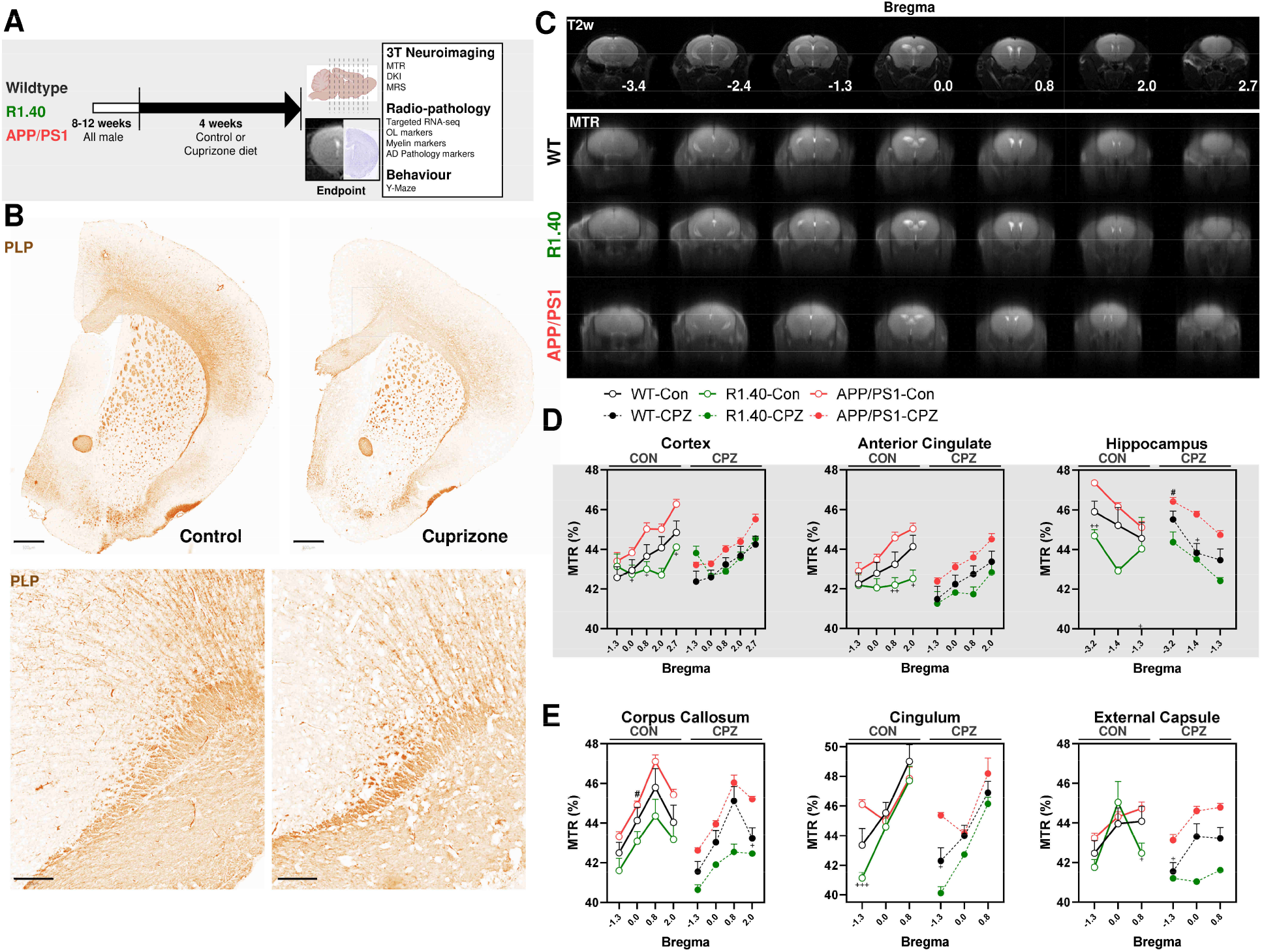
Magnetization transfer ratio (MTR) detects cuprizone-mediated demyelination in mouse models. **A**. Experimental design of the present study. Cuprizone (CPZ, 0.2% w/w diet)-mediated demyelination was conducted in WT, R1.40 and APP/PS1 male animals between 8-12 weeks old. After four weeks of CPZ, the animals were scanned by 3T MRI with magnetization transfer (MT), diffusion tensor imaging (DTI) and MR spectroscopy (MRS). The scanned brains were then collected for bregma region-specific radio-pathological investigation for myelin, oligodendrocyte and AD pathology-associated markers. The cognitive function was test by Y-Maze in a subset of animals prior to MRI scanning. **B**. CPZ-mediated demyelination for four weeks significantly reduced myelin protein expression as exemplified by PLP immunohistochemistry (Scale bar at upper panel, 500 µm, lower panel, 100 µm). **C.** Upper panel, representative T2w image of the seven coronal brain slices in MRI analysis from Bregma -3.4 (posterior) to Bregma 2.7 (anterior) in WT. Lower panel, representative image of magnetization transfer ratio (MRT) from Bregma -3.4 to 2.7 of WT, R1.40 and APP/PS1 in control condition. MTR measurements in WT (black, n = 15), R1.40 (green, n = 5) and APP/PS1 (red, n = 14) with (closed circle) or without CPZ treatment (open circle) across **D** gray matter (GM) of traced regions including cortex (Bregma -1.3 to 2.7), anterior cingulate cortex (Bregma -1.3 to 2.0), and hippocampus (Bregma -3.2 to 1.3), and **E** across white matter (WM) including corpus callosum (Bregma -1.3 to 2.0), cingulum (Bregma -1.3 to 0.8) and external capsule (Bregma -1.3 to 0.8). The statistically significant difference in each treatment group at specific Bregma region is denoted next to the symbol against WT as * or against APP/PS1 as +. The region-specific statistically significant difference between control and CPZ in each strain is denoted as # next to the symbol. Statistical analysis of between genotype or treatment in all regions performed by Two-way ANOVA repeated measures followed by Tukey’s multiple comparisons test (* or + or #, p < 0.05; +++, P<0.001)

To achieve a greater contrast, we applied magnetization transfer ratio (MTR), a sensitive MR technique for detecting myelin content (bottom rows, Fig. 1C, (68)). CPZ treatment lowered MTR in most regions of all strains, but the reduction reached significance only in the cortex (p < 0.05) and hippocampus (p < 0.05) of APP/PS1. Genotype conferred a significant difference in MTR in corpus callosum (p < 0.05), external capsule, cingulum and hippocampus. Unexpectedly, except for three specific areas (cingulum at Bregma 0.0 & 0.8 and external capsule at Bregma 0.0), R1.40 mice consistently showed the lowest MTR measurement both at baseline and after CPZ treatment (green symbols, Fig. 1D, E). These MTR results, while unexpected, suggest that the APP_SWE_ overexpression in R1.40, but not in APP/PS1, reduced the myelin content of both GM and WM across the brain and highlights R1.40 the distinct demyelination response of R1.40.

### Demyelination triggers opposite changes in R1.40 and APP/PS1 brains

To investigate if MTR findings above were the consequence of the intrinsic axonopathy reported in similar APP_SWE_ overexpression models (69–72), we applied DTI sequence to quantify the microstructural integrity of the myelinated tracts, as exemplified by the slice at Bregma 0.8 (Fig. 2A-E). The anisotropic scalar, FA, is a sensitive metric to detect the difference in directionality between the myelin tracts. As expected, we found large difference between FA values in GM (predominated by crossing fibers) and WM (predominated by fiber bundles) (Fig. 2F, G). However, neither genotype nor cuprizone contributed to significant differences in FA.

**Figure 2.**
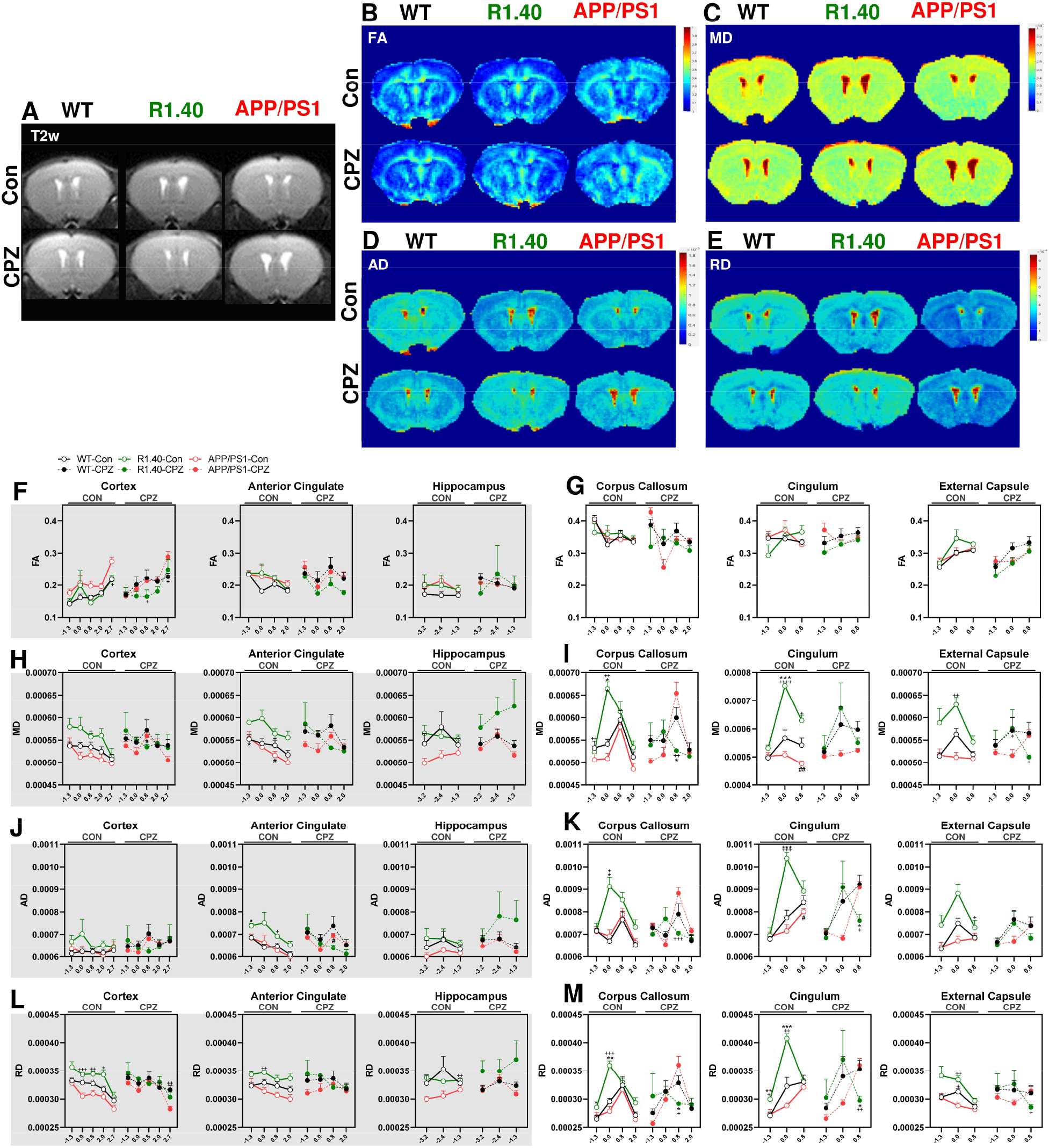
Diffusion tensor imaging (DTI) identified divergent effects of demyelination on R1.40 and APP/PS1. Representative neuroimage Bregma 0.8 of the frontal lobe for **A** T2w, **B** fractional anisotropy (FA), **C** mean diffusivity (MD), **D** Axial diffusivity (AD), and **E** radial diffusivity (RD). across WT, R1.40 and APP/PS1 in control (Con) and CPZ condition. **F-M** DTI analyses with quantifications across Bregma -3.4 to 2.7 in GM (shaded graphs; Cx, cortex; Acc, anterior cingulate cortex; hippo, hippocampus) and WM (Cc, corpus callosum; Cing, cingulum; Ec, external capsule). **F, G** In most regions, no statistical significance in FA was among genotypes or in CON versus CPZ conditions. **H, I** Significant difference in MD between genotypes was found mostly in the WM region at Bregma 0.0 where R1.40 demonstrates an opposite direction of demyelination response towards CPZ. CPZ mediated significant effects only in APP/PS1 in Cx, Acc and Cing and only at Bregma 0.8. At this slice, significant differences between R1.40 and APP/PS1 was also observed in Cx, Acc and Cing at baseline as well as Cc and Ext Cap after CPZ treatment. **J, K** The largest significant difference in axial diffusivity (AD) between genotypes was also found mostly in the WM region at Bregma 0.0 where the demyelination response in R1.40 remained opposite to other genotypes. CPZ mediated significant effects only in APP/PS1 in Acc and Cing only at Bregma 0.8. In this brain slice, a statistically significant difference between R1.40 and APP/PS1 was observed in Acc and Ext Cap at baseline as well as in Cc and Cing after CPZ. **L, M** For RD, a major difference between WT and APP/PS1 was found in Cx after CPZ. A major difference between untreated R1.40 and APP/PS1 was found in Cx between Bregma 0.0 to 2.0, as well as Acc at Bregma 0.0 and Hippo at -1.3. An opposing demyelination response was consistently observed in R1.40 with most differences between genotypes found at Bregma 0.0 and 0.8. The significantly higher diffusivity (MD, AD, RD) found in R1.40 are suggestive of an intrinsic axonopathy at baseline. The effect of genotype on microstructural changes was unexpectedly more profound than that mediated by cuprizone. The statistically significant difference in each treatment group at specific Bregma region is indicated next to the symbol against WT as * or against APP/PS1 as +. Statistical analysis of between genotype or treatment in all regions performed by Two-way ANOVA repeated measures followed by Tukey’s multiple comparisons test (* or + or #, p < 0.05; ++, P<0.01, +++, P<0.001, ++++ P<0.0001, WT, n = 16; R1.40, n = 6; APP/PS1, n = 14).

Diffusivity metrics including mean, axial and radial diffusivity (MD, AD and RD, respectively), measure the degree of water diffusion across the myelinated axons in all directions (MD), parallel to the direction of the axons the (AD) or perpendicular (radial) to them (RD). An increase of diffusivity is indicative of a microstructural change in the myelin tract. At the baseline level (CON, control; open circles), genotype was the major contributor to differences in diffusivity in every direction in the corpus callosum (MD, AD, RD, p < 0.01), external capsule (MD, AD, RD, p < 0.05), cingulum (MD, AD, RD, p < 0.001), cortex (MD, RD, p < 0.01) and anterior cingulate (MD, AD, RD, p < 0.05). No significant changes were found in hippocampus (Fig. 2H - M). CPZ demyelination increased diffusivity overall but in doing so nearly completely eliminated the difference among the genotypes (Fig. 2H-M; filled circles), with diffusivity in hippocampus being the only exception (MD, RD, p < 0.05). The untreated R1.40 mice had the highest diffusivities in all areas with few exceptions (e.g. hippocampus at Bregma -2.4). At various Bregma regions, the diffusivity level of R1.40 was significantly higher than that of APP/PS1 (as denoted by +) and/or WT (as denoted by *). This distinct genotype difference is suggestive of an inferior myelin fiber integrity in R1.40 animals.

In keeping with these strain differences, demyelination led to opposing diffusivity responses in APP/PS1 and R1.40 mice (Fig. 2H - M). Whereas cuprizone-mediated demyelination led to an increase of diffusivities in APP/PS1 and, to a lesser extent, in WT animals, demyelination decreased diffusivity in R1.40 mice. This effect was most notable in the cortex, anterior cingulate, corpus callosum, cingulum and external capsule, particularly at Bregma 0.8. In this area, GM (cortex, anterior cingulate cortex) and WM (corpus callosum, cingulum, external capsule) regions were affected by both genotype and cuprizone. These observations agree with the MTR results and suggest that both the myelin content and microstructural integrity are compromised by APP_SWE_ overexpression in R1.40, but not in the genetically more complex APP/PS1.

### Demyelination initiates a metabolic shift in R1.40 but not APP/PS1 or WT

To investigate the neurochemistry underlying the MRI findings, an additional voxel encompassing cortex, corpus callosum, and hippocampus between Bregma -1.3 to -3.2 was analyzed for DTI changes and MRS measurements (Fig. 3A). Consistent with the observations above, no difference in FA was identified across mouse strains or between untreated and cuprizone-treated subjects. Genotype led to a significant difference in MD (p = 0.017) and RD (p = 0.0044) across brain slices (Fig. 3B). Again, R1.40 stood out from both other genotypes as having the greatest diffusivities (MD, AD and RD) before treatment, and CPZ treatment led to an obvious trend for demyelinating response in R1.40 and APP/PS1 to move in opposite directions. Thus, across most regions, the diffusivities decreased in R1.40 but increased in APP/PS1 (Fig. 3C).

**Figure 3.**
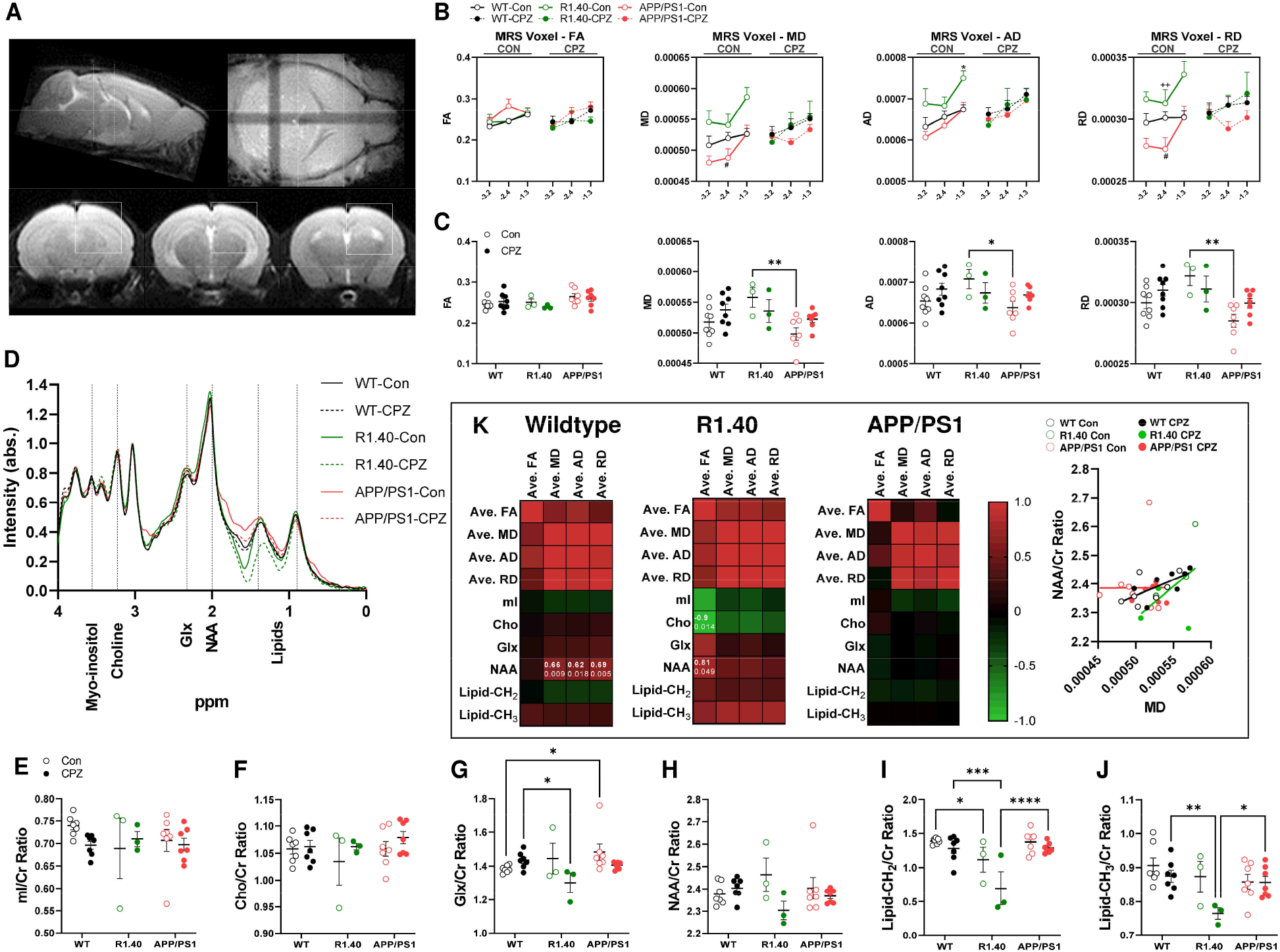
Magnetic resonance spectroscopy (MRS) confirms differential demyelination response in WT, R1.40 and APP/PS1. **A** Representative sagittal, axial and coronal T2w image of the voxel specified for MRS analysis at 3T. The 3 × 3 × 3 voxel encompasses cortex, hippocampus and corpus callosum from Bregma -3.2 to -1.3. **B** DTI analysis of this voxel confirms the opposing demyelination response in R1.40 animals across MD, AD and RD measurement with no differences in FA. **C** The average measurement of DTI finding across the voxel found significantly higher MD, AD and RD between R1.40 and APP/PS1, but not between WT and APP/PS1 (Two-way ANOVA repeated measures followed by Tukey’s multiple comparisons test, (*p < 0.05; **P<0.01). **D** The MRS spectra plot (0 – 4 ppm) with curves for WT (black), R1.40 (green) and APP/PS1 (red) in control (solid line) or CPZ treatment (dash line) with chemical shifts denoted on x-axis. The peak at **E** 3.56 ppm for myo-inositol, **F** 3.23 ppm for choline, **G** 2.3 ppm for Glx (glutamine + glutamate), **H** 2.0 ppm for N-acetyl aspartate (NAA), **I** 0.9 ppm for CH3-Lipid and **J** 1.3 ppm for CH2-Lipid were compared between all groups, where Glx/Cr, NAA/Cr, CH2-Lipid/Cr and CH3-Lipid/Cr were significantly lowered in R1.40 than APP/PS1 and WT after CPZ. Two-way ANOVA repeated measures followed by Tukey’s multiple comparisons test, (*p < 0.05; **P<0.01) **K** Pearson correlation matrix between DTI and MRS findings in this voxel showing a distinct pattern of correlation in WT, R1.40 and APP/PS1, where Glx and CH3-Lipid significantly correlated with MD, AD and RD in WT only. (All based on Pearson test with r value and p value indicated within the square with statistical significance. WT, n = 15; R1.40, n = 6; APP/PS1, n = 14). The correlation between NAA/Cr and MD is exemplified by the X-Y plot on the right.

MRS measurements showed distinct spectra in WT, R1.40 and APP/PS1 with or without demyelination (Fig. 3D). We paid particular attention to myo-inositol (peak at 3.56 ppm), choline (3.23 ppm), Glx (glutamine + glutamate, 2.3 ppm), NAA (N-acetyl aspartate, 2.0 ppm), methyl-group containing lipid/macromolecule (CH_3_-Lipid-, 0.9 ppm) and methylene-group containing lipid/macromolecule (CH_2_-Lipid, 1.3 ppm) normalized to the level of creatine (Cr; Fig. 3E-J). No significant difference was found in myo-inositol/Cr and choline/Cr ratios (Fig. 3E, F). While genotype and cuprizone did not lead to a significant change, the lowest level Glx/Cr and NAA/creatine was found in R1.40 animals after demyelination. Both Glx/Cr and NAA/Cr were lower in R1.40 than WT after cuprizone treatment (-9.3%, p = 0.0257; -4.2%, p = 0.077, respectively, Fig. 3G, H). The largest changes were in lipid and macromolecules, where both genotype and CPZ contributed to a significant change of CH_2_-Lipid (p = 0.0001, p = 0.006) and CH_3_-Lipid (p = 0.03, p = 0.02), respectively. Similarly, the lowest level of lipid/Cr was observed in R1.40 upon demyelination (CH_2_-Lipid % -45.6% vs WT, -45.9% vs APP/PS1; CH_3_-Lipid % -12.7% vs WT, -10.8% vs APP/PS1), as shown in Fig. 3I, J.

To provide a better understanding of the relationship between diffusivity and these chemical shifts, we performed Pearson correlation between DTI and MRS for WT, R1.40, and APP/PS1 scans. In WT, positive correlations were only significant between diffusivity scalers (MD, AD, RD) and NAA/Cr (Fig. 3K, heatmap). The strong positive correlation between NAA/Cr and MD in WT reflects the fact that NAA is a metabolite biomarker for compromised myelin tract integrity (Pearson r = 0.665, p = 0.0095, Fig. 3K, X-Y plot). In contrast, R1.40 and APP/PS1 demonstrated a distinct pattern of correlation between diffusivity and MRS finding with no statistical significance. The consistently low level of Glx/Cr, NAA/Cr and Lipid/Cr in R1.40 upon demyelination indicates change of brain chemistry that could be a result of diffuse axonal damage and/or metabolic shift (73, 74). Overall the patterns suggest that there are differential responses towards demyelination among the three strains.

### Metabolic shift, but not axonal changes in R1.40 and APP/PS1

R1.40 is known to express less Aβ than APP/PS1 animals across the lifespan (39). The neuroimaging results therefore disagreed with our first hypothesis that more severe amyloid pathology leads to more severe myelin pathology. To determine if pre-existing axonal changes in R1.40 might explain the neuroimaging findings (69–72), we examined the expression of pan-neurofilament or βIII-tubulin in the cerebral cortex. Western blots revealed that neither genotype nor drug exerted any significant effects on the two protein (Fig. 4A, B). βIII-tubulin immunohistochemistry of the frontal cortex confirmed these findings (Fig. 4C). These data, therefore, suggest that there are no major axonal deficits at these early ages in R1.40 animals. No observable difference in the measurement of βIII-tubulin-IR or semi-quantification of βIII-tubulin^+^ axon density was found across WT, R1.40 and APP/PS1 with or without cuprizone treatment (Fig. 4C).

**Figure 4.**
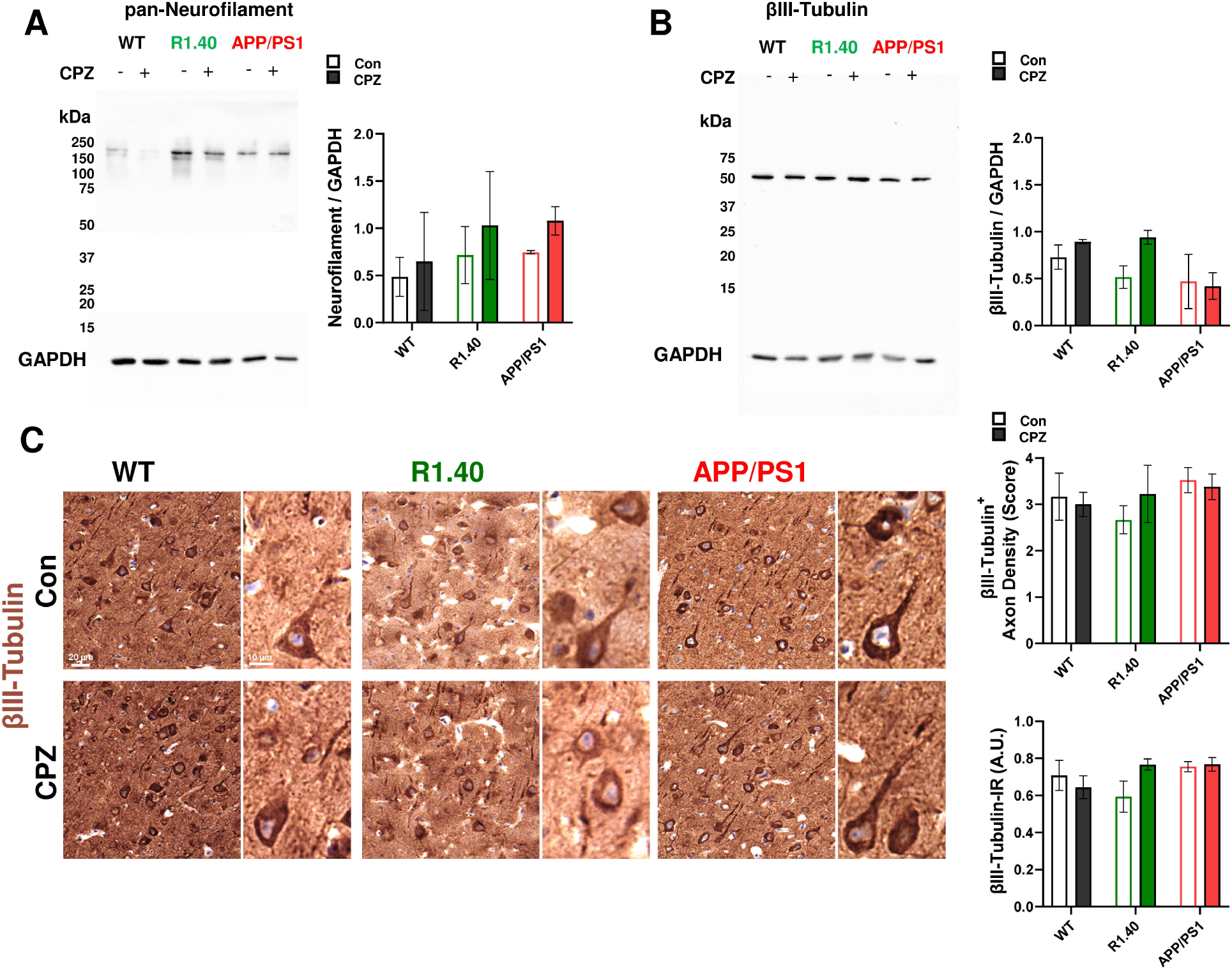
Axonal pathology was minimal across the cerebral cortex in WT, R1.40 and APP/PS1. The effects of genotype and demyelination on the axonopathy in cortex were tested by immunoblotting experiments. Representative blot of pan-neurofilament and βIII-tubulin, two specific markers of axon-specific cytoskeleton, in the mouse cerebral cortices were shown in **A** and **B**, respectively. No significant difference was observed between WT, R1.40 and APP/PS1 animals with a slight increase upon demyelination only in R1.40 (n = 3). GAPDH is used as the loading control. **C** *Left* Representative immunohistochemistry image of βIII-tubulin on coronal brain section at MCx of Bregma 0.8 in WT, R1.40 and APP/PS1 with or without CPZ treatment. No significant difference was observed. *Right* The density of βIII-tubulin+ axons were graded by 3 independent experimenters on randomized images at x20 (WT n = 13, R1.40 n = 7, APP/PS1 n = 14) and no difference was detected. Measurements of βIII-tubulin-IR signal in the same region also showed no differences. Statistical analysis of between genotype or treatment in all regions performed by Two-way ANOVA followed by Tukey’s multiple comparisons test.

We next examined the transcription profile of cells in the cerebral cortex with targeted RNA-seq (Fig. 5A). The RNAs from the WT, R1.40 and APP/PS1 cortex with or without cuprizone treatment were subjected to analysis using a panel of the 416 genes covering nine major pathways associated with age-related pathology, including angiogenesis (12.2%), apoptosis (8.5%), cell cycle (18.3%), cell senescence (6.3%), DNA damage & repair (14.8%), epithelial-to-mesenchymal transition (11.1%), hypoxia (12.2%), metabolism (11.1%) and telomeres/telomerase (5.7%, see gene list in Table S1).

**Figure 5.**
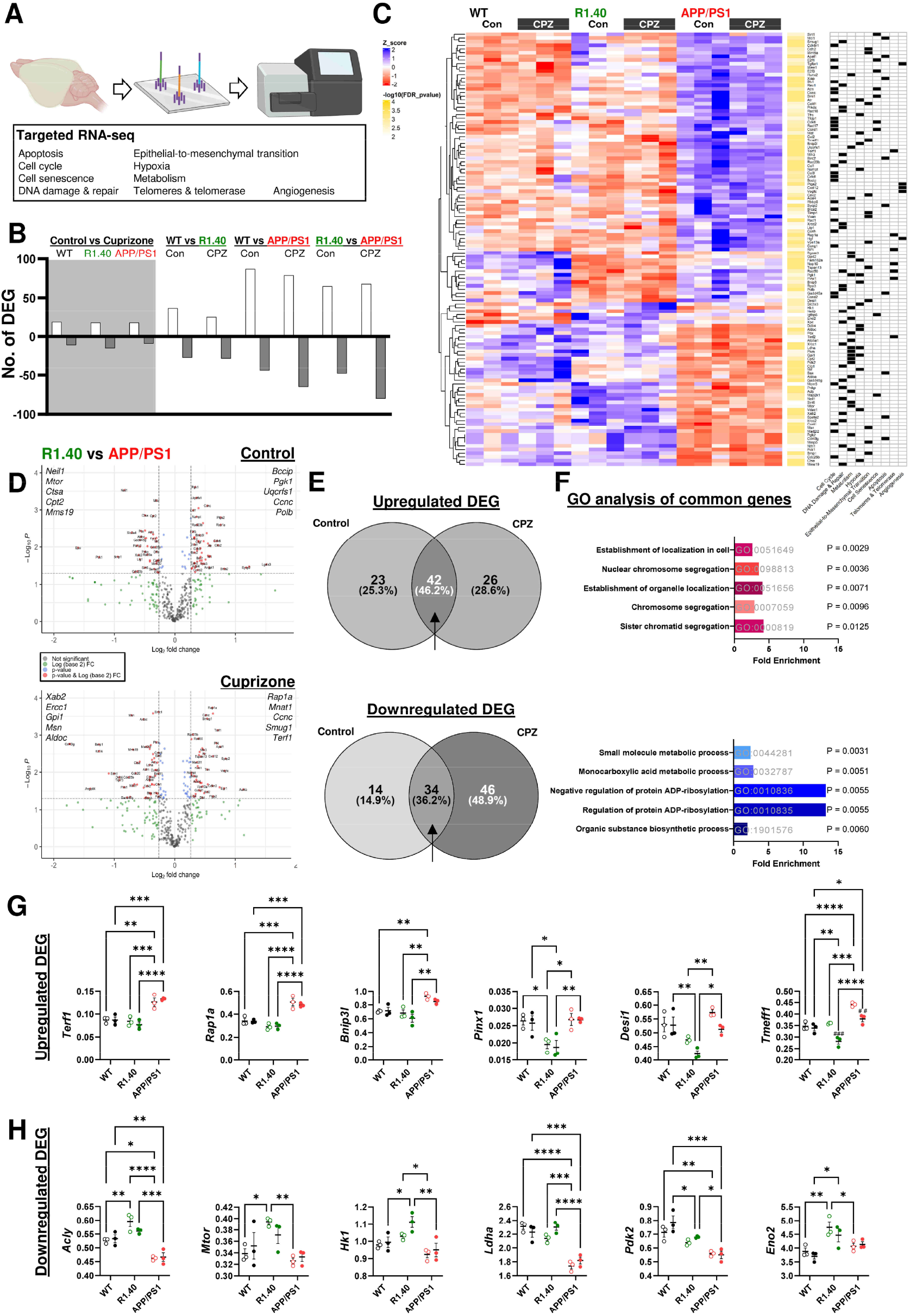
Differential gene expression in WT, R1.40 and APP/PS1 at baseline and after cuprizone treatment. **A** Cerebral cortex of WT, R1.40 and APP/PS1 animals with or without cuprizone treatment was isolated for RNA extraction and a targeted RNA-seq transcriptome panel targeting 416 genes covering nine pathways associated with aging on a MiSeq system. **B** The number of significant differentially expressed genes (DEGs, upregulated or downregulated) were much higher between genotype pairs (WT vs R1.40, WT vs APP/PS1, R1.40 vs APP/PS1) than that between treatments (Control vs cuprizone). **C** Heat map showing the unique gene signature among the 119 DEGs across the three mouse strains with or without cuprizone treatment. **D** Volcano plot showing the and DEGs between R1.40 and APP/PS1 in control and cuprizone with the top upregulated and downregulated genes stated. E Comparison between the DEGs in **D** identified 42 and 34 genes commonly upregulated and downregulated in APP/PS1 with or without cuprizone treatment (arrow). **F** Gene ontology analysis with PANTHER-based enrichment showing the top five biological processes significantly enriched in the DEGs commonly upregulated or downregulated in APP/PS1. The GO term references and the enrichment significance are denoted on the graph. The representative transcript expression in the top enriched processes from, **G** upregulated DEGs (*Establishment of localization in cell*, GO:0051649) and, **H** downregulated DEGs (*Small molecule metabolic process*, GO:0044281). The significant difference between genotype are denoted by two-way ANOVA among with Tukey’s multiple comparisons test, * p < 0.05, ** p < 0.01, *** p < 0.001, **** p <0.0001; n = 3 per group, n = number of independent animal subject. Of note, *Rap1a*, *Bnip3l*, *Pinx1*, *Desi1* and *Tmeff1* are known to be highly expressed by OL or OPC in normal mouse cortex according to single cell RNA database, DropViz (http://dropviz.org/), as shown in Fig. S1D, E (75).

Examination of the differential expressed genes (DEGs) revealed that by far the largest effect was due to genotype with WT, R1.40 and APP/PS1 differing markedly from one another. By contrast, cuprizone induced a significant change in only 6-8% of the transcripts, regardless of genotype (Fig. 5B). Note that the transcription profile of APP/PS1 differed substantially from WT and R1.40 (Table S1 and PCA in Fig. S1). Genotype contributed to a significant difference in 119 genes (FDR < 0.01) across strains as illustrated in Fig. 5C. These genes were distributed across the nine pathways. Consistent with the neuroimaging, a major difference was found between R1.40 and APP/PS1 (Fig. 5D). The commonly upregulated (46.2%) and downregulated (36.2%) genes between R1.40 and APP/PS1 in control and cuprizone were identified and a GO analysis against the panel background was performed (Fig. 5E, F). A significant enrichment in *establishment of localization in cell* (GO:0051649, Fold = 2.59, p = 0.0029) and *small molecule metabolic process* (GO:0044281, Fold = 2.46, p = 0.003) were identified among the top up- and down-regulated genes, respectively (Fig. 5F).

Our findings align with previous single cell RNA-seq as we found *Rap1a*, *Bnip3l*, *Pinx1*, *Desi1* and *Tmeff1* to be significantly upregulated in APP/PS1. All are significantly enriched in the OL/OPC population of the mouse frontal cortex (Dropviz (75), Fig. S1). Importantly, transcripts significantly upregulated only in R1.40 such as *Acly*, *Mtor* and *Hk1* (Fig. 5H) are important to axonal transport, autophagy, and glucose metabolism (76). This is consistent with our MRS findings where a major difference in brain metabolism exists between R1.40 and APP/PS1.

### APP/PS1 mice express a higher level of human APP protein than R1.40 mice in the cerebral cortex

As few if any amyloid plaques have formed, the phenotype of R1.40 and APP/PS1 are rarely investigated at young ages (e.g. < 4 months) (38, 39, 77). We investigated the expression of APP protein in the cerebral cortex with the 6E10 antibody that recognizes residues 1-16 of the human β-amyloid peptide, and the Y188 antibody that recognize the C-terminus of both human and mouse amyloid precursor protein (clone Y188, amino acids 750-770) (Fig. 6). Immunoblots probed with 6E10 revealed that levels of APP were 1.8 times higher in APP/PS1 than R1.40 brain (p = 0.043) and 6.6 times higher than WT (p = 0.0004). These differences were maintained after cuprizone treatment. As expected, at these young ages, no 6E10 Aβ peptide was detected despite prolonged exposure. Similarly, APP protein expression detected by Y188 antibody was significantly higher in APP/PS1 than in WT by 2.49-fold (p = 0.0494, Fig. 6B). As the Y188 antibody will also detect the cleaved APP C-terminal fragments (CTFα and CTFβ), we used longer exposures to reveal CTFs in both R1.40 and APP/PS1. The variability was high, but we found no significant changes of CTF with cuprizone treatment in either transgenic line (Fig. 6C). These results indicate that APP/PS1 brain express a higher level of human APP than R1.40, and that *y*-secretase processing of the full-length APP protein is equivalent even though the APP/PS1 mice carry an extra PSEN1-encoding transgene. Notably, cuprizone did not exert any significant effects on the overall APP protein expression in any genotype strains.

**Figure 6.**
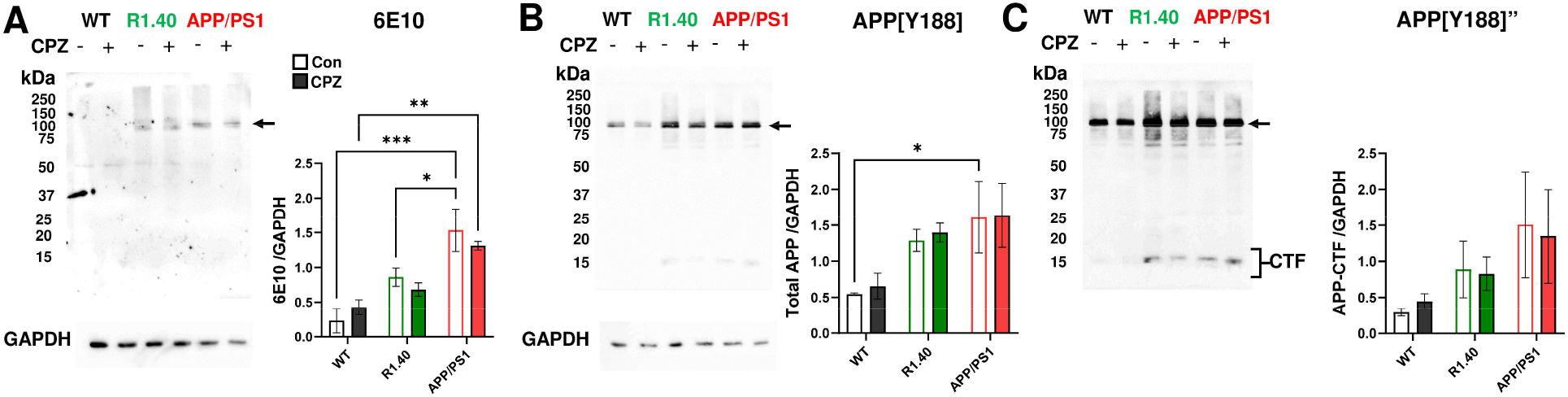
Amyloid precursor protein expression is the strongest in APP/PS1 cortex lysate. Representative immunoblots of antibody against **A** β-amyloid (6E10) of human origin and, **B** amyloid precursor protein c-terminal residues a.a 750-770 (clone Y188) of human and mouse origin. **A** Quantifications showed that the strongest 6E10 signal at ∼110 kDa representing whole human APP (∼110 kDa) was detected in APP/PS1. No soluble β-amyloid from 4 to 20 kDa was detected. No 6E10 expression was detected in WT. **B** Similarly, the strongest Y188 signal at ∼110 kDa showed the strongest whole APP (∼110 kDa) in APP/PS1. **C** APP[Y188]” is the blot with longer exposure to visualize the cleavage c-terminal fragments (CTF) of APP as found in R1.40 and APP/PS1 (∼15-25 kDa), which is indicative of active APP processing. Although the strongest CTF was found in APP/PS1, no statistical significant difference was observed. In all blots, cuprizone-mediated demyelination had no effects on the protein expression level of APP in any strains. Statistical analysis of between genotype or treatment in all regions performed by Two-way ANOVA followed by Tukey’s multiple comparisons test. * p < 0.05, ** p < 0.01, *** p < 0.001, **** p <0.0001; n = 3 per group, n = number of independent animal subject.

### Demyelination increase 6E10 immunoreactivity only in APP/PS1

The distribution of amyloid deposition across the cerebral cortex is unequal in APP/PS1 and other AD transgenic mice (78). To ask if cuprizone-mediated demyelination accelerated amyloid pathology in a region-specific way, we used immunohistochemistry to track the pattern of amyloid proteins in frontal cortex (Bregma 0.8), where a major difference in DTI was found between R1.40 and APP/PS1 (Fig. 2) and where intracortical myelination is strongly associated with cognitive functions (14, 15). At the macroscopic level, the expression pattern of human APP detected by 6E10 antibody was consistent with the immunoblotting results. APP/PS1 animals showed the strongest immunoreactivity (-IR) followed by R1.40 with very low level in WT (Fig 7A). Importantly, 6E10-IR was significantly upregulated by cuprizone only in APP/PS1 mice (p < 0.0001). With particularly strong effect across the Acc, (anterior cingulate cortex), MCx (primary motor cortex), and SSCx (somatosensory cortex), 6E10-IR was increased 51.1% in the frontal cortex of APP/PS1 (p = 0.0005, Fig. 7B). As suggested by the absence of low molecular weight Aβ in brain lysate, no diffuse amyloid plaques were detected in any brains examined.

**Figure 7.**
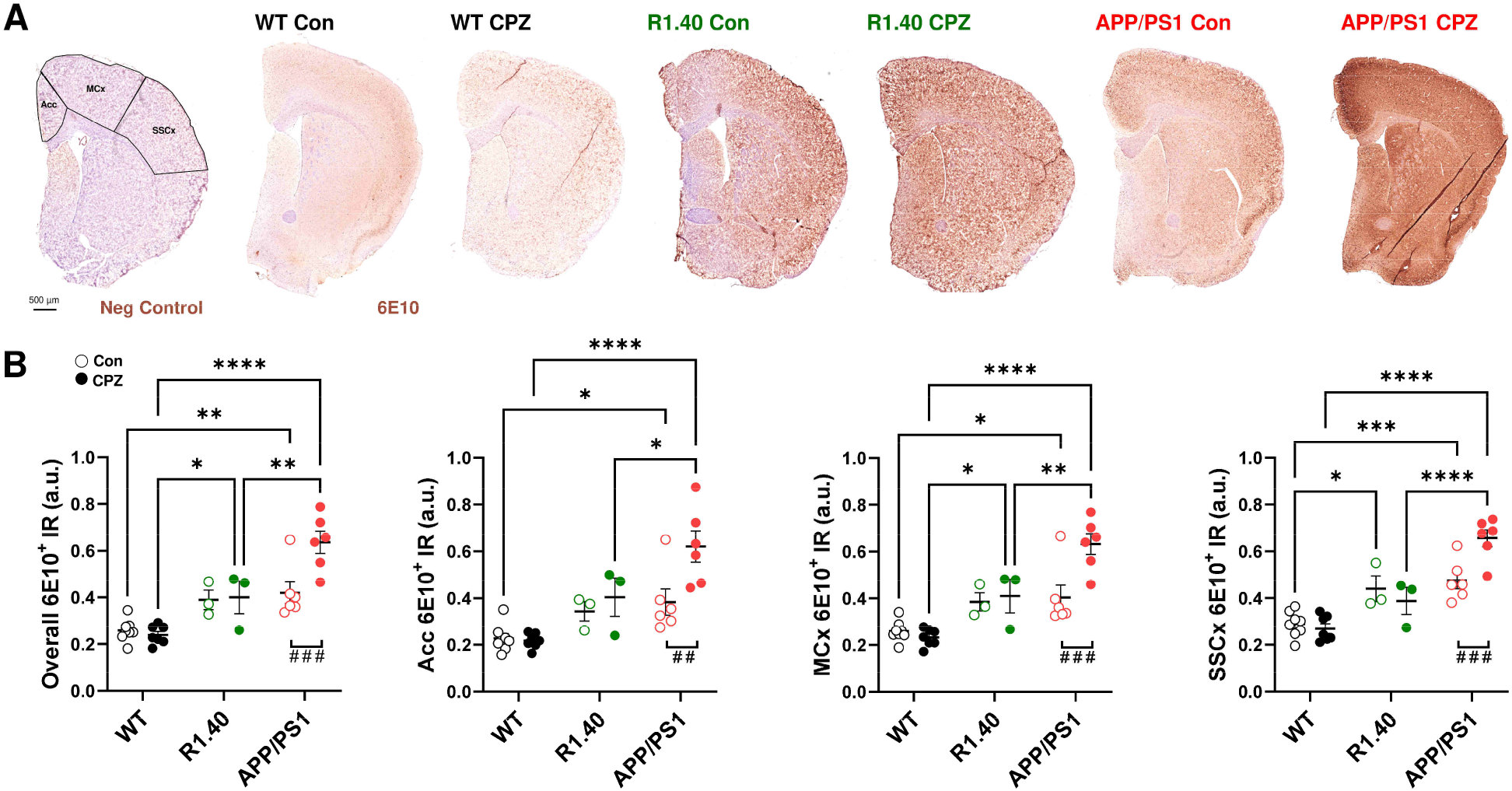
Cuprizone-induced acute demyelination increased 6E10-IR only in APP/PS1 mice but not R1.40. **A** Representative whole slide image of 6E10 (β-amyloid) immunoreactivity(-IR) on coronal brain section at Bregma 0.8 in WT, R1.40 and APP/PS1 with or without CPZ treatment. 6E10-IR across Acc, MCx and SSCx were measured as indicated by the annotation on the negative control. The strongest 6E10 signal was observed in APP/PS1. **B** Quantification of 6E10-IR showed a significantly higher signal in R1.40 and APP/PS1 compared with WT in the average cortex measurement (overall) and in each region. CPZ treatment significantly increased 6E10-IR only in APP/PS1 across each cortical regions. Statistical analysis performed by Two-way ANOVA followed by Tukey’s multiple comparisons test (n = 3 – 7, versus WT, *p < 0.05; **p < 0.01, ****p < 0.0001; versus CPZ, ##p < 0.01, ###p < 0.001). Acc, anterior cingulate cortex; MCx, primary motor cortex; and SSCx, somatosensory cortex

### Cuprizone triggers a widespread intraneuronal amyloid deposition across the brain in APP/PS1

Given the unexpected and distinct differences between R1.40 and APP/PS1 in neuroimaging, transcription and amyloid response to demyelination, we focused solely on APP/PS1 for all subsequent experiments. As demyelination triggered an increase of 6E10 immunostaining in APP/PS1 mice, we asked whether accelerated amyloid pathology could be found in regions beyond the cortex. We examined coronal sections of the brain regions most vulnerable to demyelination as defined by our MRI findings (Bregma 0.8 and Bregma -1.3, see DTI). Immunohistochemistry of 6E10 was examined across cerebral cortices, basal nucleus, and thalamic regions (Fig. 8A). We found that cuprizone treatment led to a significant increase in the overall level of 6E10-immunoreactvity(-IR) level as well as an increase in the density of cells 6E10^+^ cells in the soma and proximal dendrites and axon-resembling regions. The CPZ effect was seen in both WT and APP/PS1 animals and was most dramatic in anterior regions especially motor cortex, somatosensory cortex and caudoputamen across Bregma 0.8, and smaller but still significant effect in the anterior cingulate cortex (Fig. 8C, D). Despite the increase of 6E10-IR, no diffuse plaques were found in any regions. Cuprizone also increased 6E10-IR in the white matter tracts of the basal ganglia as well as in the cingulum/corpus callosum of APP/PS1 (Fig. 8A, CP, Cg/CC). This suggests that demyelination induced strong APP expression along the axons of the major projection pathways.

**Figure 8.**
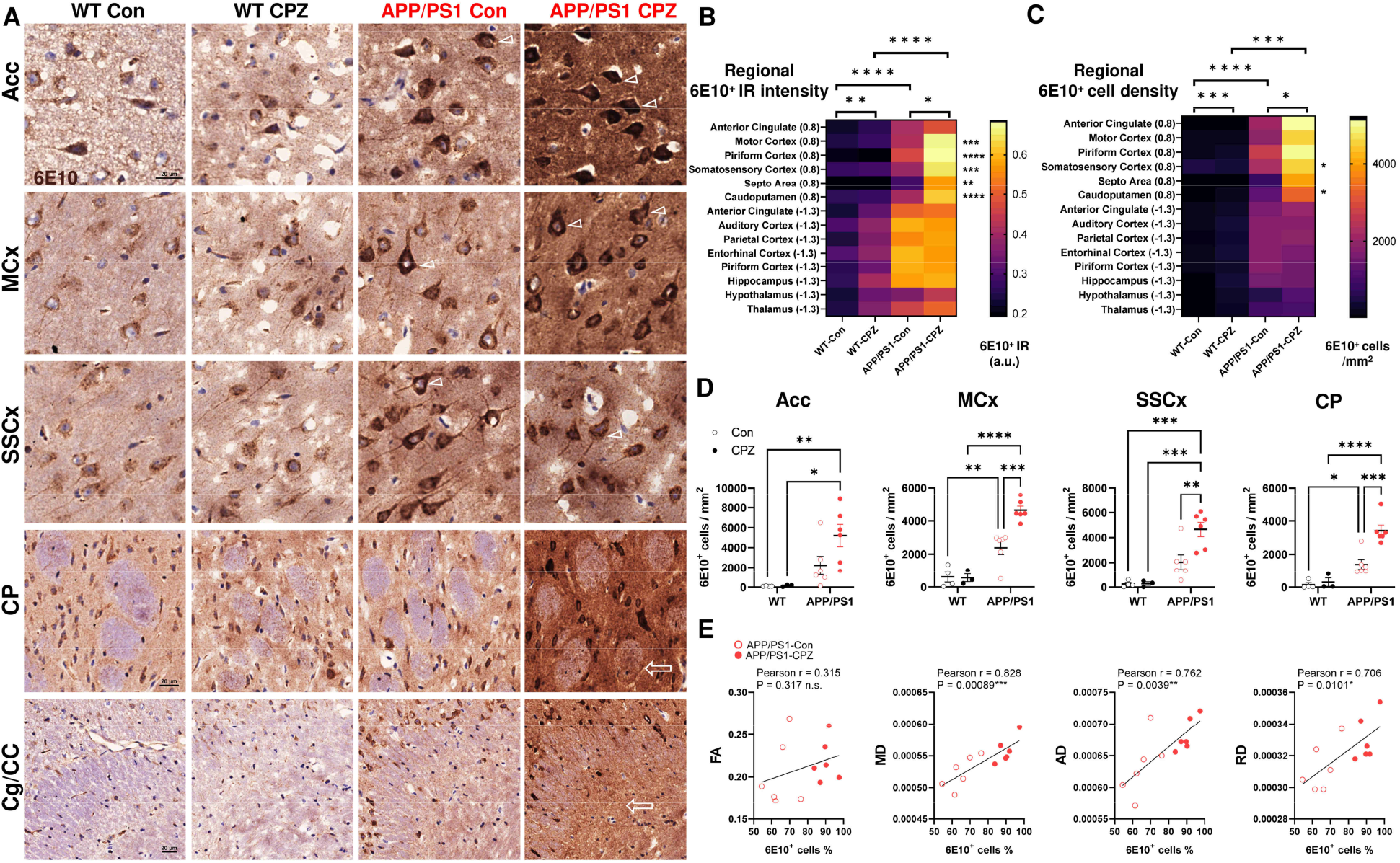
Demyelination exacerbated region-specific intraneuronal amyloid deposition in APP/PS1. **A** Representative microscopic image of 6E10 (β amyloid) immunoreactivity(-IR) on across Acc, (anterior cingulate cortex), MCx (primary motor cortex), SSCx (somatosensory cortex), CP (Caudoputamen) and Cg/CC (cingulum and corpus callosum) at Bregma 0.8 in WT and APP/PS1 with or without CPZ treatment. A genotype- and cuprizone-dependent increase of 6E10-IR was apparent within the neurons (open arrowheads) and across the neuropil in the GM. Most of the 6E10 signals were within neuronal soma and suggestive of the accumulation of intraneuronal amyloid. These increases of 6E10 signal were also found within the myelinated bundles in CP and Cg/CC (open arrows). Yet, no diffuse amyloid plaques were found in any brain regions. **B** Quantification of 6E10-IR (left) and 6E10^+^ cells density at Bregma 0.8 and -1.3 across eleven brain structures revealed a significant genotype- and CPZ-dependent the 6E10. **C, D** The CPZ-driven, region-specific increase of 6E10-IR and 6E10^+^ cells density at Acc, MCx, SSCx and CP. **E** The percentage of 6E10^+^ cells across the MRI-scanned APP/PS1 cortex at Bregma 0.8 was plotted against the DTI measurements (FA, MD, AD and RD). Strong (r > 0.7) and statistically significant correlations were found between MD, AD and RD against 6E10^+^ cells, but not FA. Statistical analysis of between genotype or treatment in all regions performed by Two-way ANOVA followed by Tukey’s multiple comparisons test (n = 4 – 6, *p < 0.05; **p < 0.01, ***p < 0.001, ****p < 0.0001). Correlation analyzed by Pearson test with r and P values denoted on graph.

To determine whether this buildup of intraneuronal amyloid deposition played a role in MRI-detected changes, we performed a subject-by-subject correlation of MRI measurements and the percentage of 6E10^+^ cells across the MRI-scanned cortex (at Bregma 0.8). We found that intraneuronal 6E10-IR was positively correlated with diffusivity scalars (MD, AD, RD), but not with FA, in APP/PS1 (Fig. 8E). No correlation was found in the WT, and no correlation between OL density and intraneuronal amyloid deposition was found in APP/PS1 (data not shown). The correlation between diffusivity and intraneuronal amyloid after demyelination in APP/PS1 mice suggests that the observed neuronal amyloidosis may be a secondary consequence of the induced myelin abnormalities.

### Short-term cuprizone has no effects on hippocampus and cognitive function

We next investigated the effect of cuprizone in the hippocampus, a highly vulnerable region in AD. Although we found a significantly higher 6E10-IR across the tissue and a greater density of 6E10 immunopositive cell in APP/PS1, cuprizone-mediated demyelination had no effects on intraneuronal amyloid deposition (Fig. 9A, B). Unlike the frontal cortex, the percentage of 6E10^+^ cells in the hippocampus (Bregma -1.3) showed no correlations with any diffusion tensor measurements (Fig. 9C). The lack of cuprizone-mediated effect in hippocampus was consistent with the absence of cuprizone-induced memory loss. In a two-tiered Y-Maze behavioral test for spatial memory (Fig 9D, E), no significant reduction of spontaneous alteration was found (Upper row, Fig. 9D). In the spatial reference memory test, no statistical difference was found in either the duration or the number of entries that the animals made into the correct arm (Lower row, Fig. 9D). Although APP/PS1 mice showed a consistently higher speed of movement, there were no changes in performance in either WT or APP/PS1 mice after cuprizone treatment. The distribution of 6E10^+^ intraneuronal amyloid deposition in these 3-month-old animals after four weeks of treatment is consistent with the reported spread of amyloid pathology (from isocortex, to entorhinal cortex to hippocampus) reported in the older APP/PS1 animals (78).

**Figure 9.**
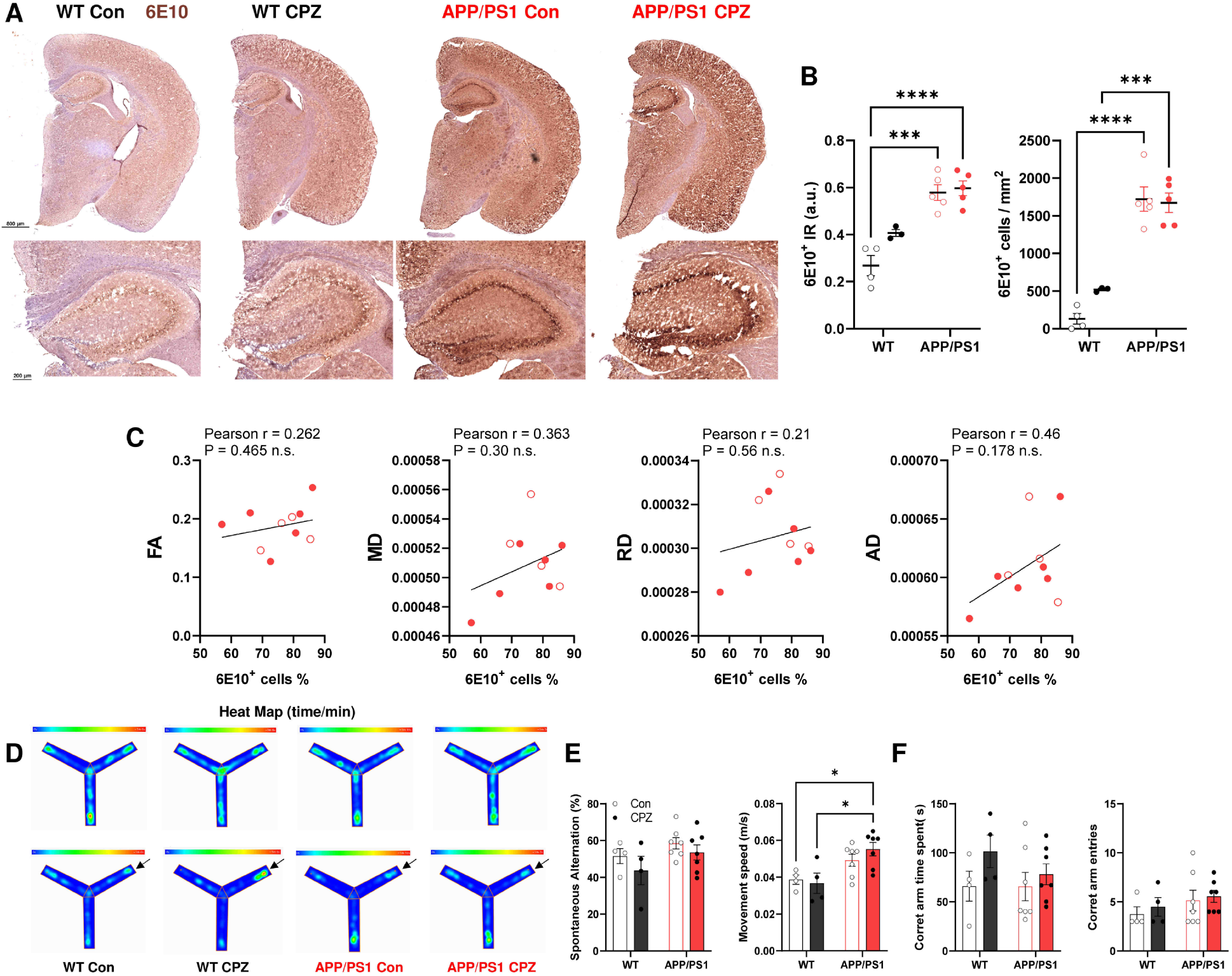
Cuprizone-induced acute demyelination had minimal effects on spatial memory and hippocampal pathology. **A** Representative whole slide image of 6E10 (β-amyloid) immunoreactivity(-IR) on coronal brain section at Bregma -1.3 in WT and APP/PS1 with or without CPZ treatment (*Above*). The corresponding 6E10-IR changes in the hippocampus at a higher magnification is shown (*below*). **B** Quantification showed the 6E10-IR and 6E10^+^ cell density in the hippocampus was significantly higher in APP/PS1 than in WT, but CPZ showed no significant effects. **C** Unlike the cortex, the percentage of 6E10^+^ cells across the MRI-scanned APP/PS1 hippocampus at Bregma -1.3 was plotted against the DTI measurements (FA, MD, AD and RD). No statistically significant correlations were found. **D** Representative heatmap of movements in Y-Maze behaviour test for spatial working memory (*Above*) and reference memory (*Below*, with arrow indicating correct arm entries). **E** No significant difference in spontaneous alternation percentage between genotype or CPZ was detected. A significantly faster speed of movement was detected in APP/PS1 mice in spatial working memory test. **F** No significant difference in time spent nor entry to the correct arm in the spatial reference memory test was found. Statistical analysis of between genotype or treatment in all regions performed by Two-way ANOVA followed by Tukey’s multiple comparisons test (n = 4 – 6, *p < 0.05; **p < 0.01, ***p < 0.001, ****p < 0.0001). Correlation analyzed by Pearson test with r and P values denoted on graph.

### APP_SWE_/PSEN1_dE9_ transgene renders oligodendrocytes more vulnerable

Our neuroimaging results demonstrated that cuprizone increased intraneuronal amyloid deposition only in the APP/PS1 animals. Previously, demyelination was found at 6 months old of age in another double transgenic AD model (with a mutant human APP_SWE_ cDNA and a mutant PS1_L166P_, both under *Thy1* promoter) (10). To ask which came first, myelin loss or amyloid pathology, we asked whether the APP_SWE_/PSEN1_dE9_ transgene exerted any deleterious effects on OL lineage at 3 months of age, before any diffuse plaques are found. We found that cuprizone-mediated demyelination triggered the loss of PLP in WT and APP/PS1 across GM and WM (Fig. 10A), in a pattern that was distinct from abaxonal myelin protein MOG (Fig. 10B). Using OL differentiation markers including Olig2 (pan-OL), Nkx 2.2 (pre-OL/committed oligodendrocyte progenitor cells, OPC), ASPA (mature OL, [mOL]) and CC1 (mOL), we further mapped and analyzed the OL lineage density in both GM (somatosensory cortex, primary motor cortex and anterior cingulate cortex) and WM (corpus callosum, external capsule and cingulum) at Bregma 0.8 and Bregma -1.3 (Fig. 10C-F). At this young age, the effect of genotype was most pronounced in the GM (Olig2^+^ and ASPA^+^ in the cortex) with virtually no area of WM showing a significant difference in APP/PS1 mice. The OL lineage in both regions responded to cuprizone and the most significant response was in the mature cells of the OL lineage (ASPA^+^ & CC1^+^). We also found distinct GM/WM differences in cuprizone response. The GM OL in APP/PS1 were far more vulnerable than that in WT animals. In the WM, on the contrary, the cuprizone effects were nearly equivalent between the two genotypes. We note the in the WT WM, cuprizone also appeared to affect the pre-myelinating OL (Nkx 2.2^+^) but not in APP/PS1. In the hippocampus, consistent with lack of intraneuronal amyloid deposition, there was no significant difference in the Olig2^+^, Nkx 2.2^+^, ASPA^+^ and CC1^+^ OL population among all groups (Fig. S2).

**Figure 10.**
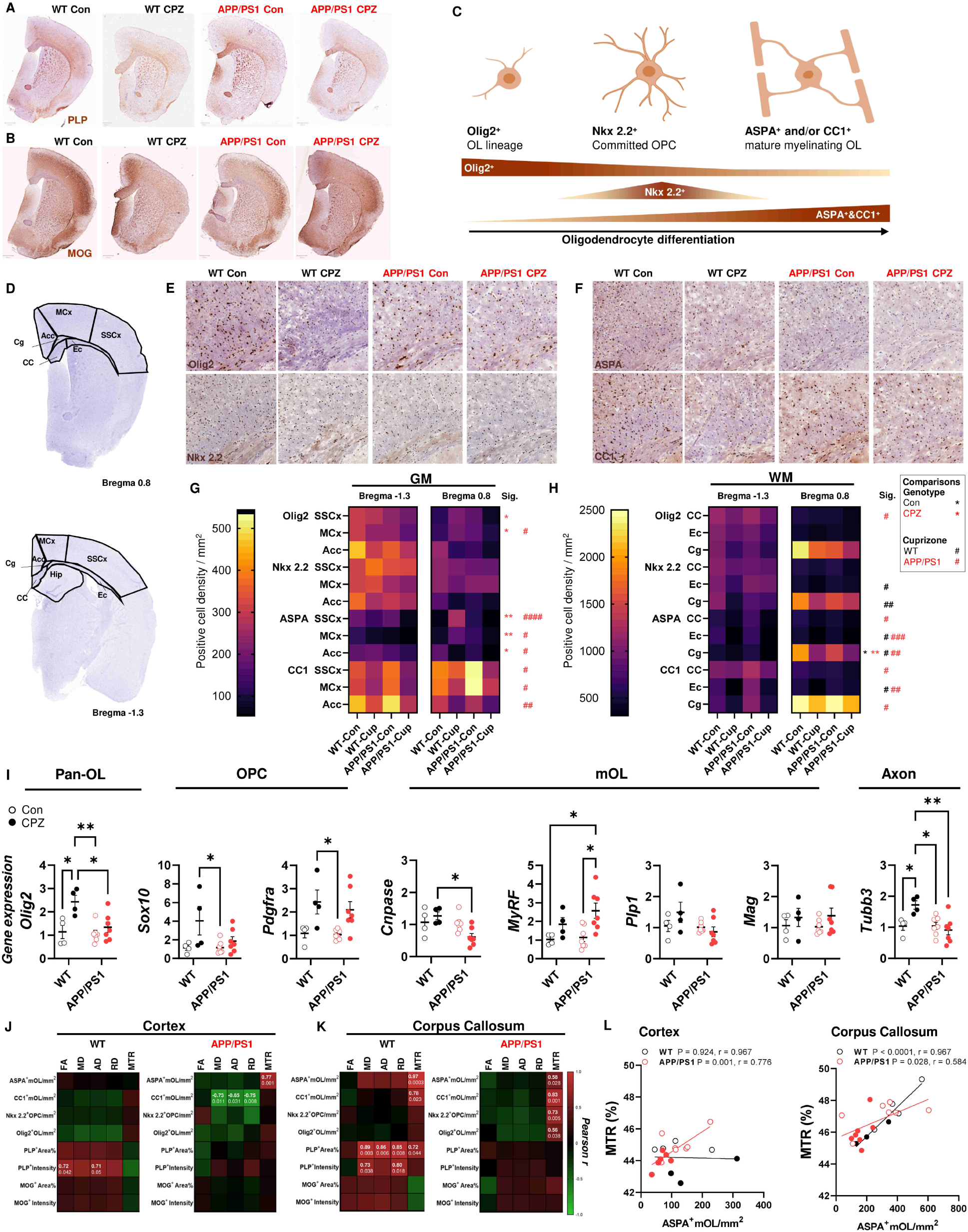
Mature oligodendrocyte in the gray matter is highly vulnerable to demyelination in APP/PS1. Representative whole slide images of **A** PLP and **B** MOG immunoreactivity(-IR) on coronal brain section at Bregma 0.8 in WT and APP/PS1 with or without CPZ treatment. The PLP-IR and MOG-IR showed distinct patten before and after CPZ. **C** Schematic diagram of OL lineage differentiation and the OL markers employed in the histology experiment where Olig2 served as a pan-OL marker, Nkx 2.2 as pre-OL/committed OPC marker while ASPA and CC1 as mOL markers. **D** Annotation maps of regions quantified for OL markers including GM regions (Acc, anterior cingulate cortex; MCx, primary motor cortex; SSCx, somatosensory cortex and Hippo, hippocampus) and WM regions (Cg, cingulum; CC, corpus callosum and Ec, external capsule) on brain slices at Bregma 0.8 and -1.3. **E, F** Representative immunohistochemistry images showing Cg/CC region with the immunoreactivity (brown, DAB-IR; blue, nuclear hematoxylin) of Olig2, Nkx 2.2, ASPA and CC1 in WT and APP/PS1 with or without CPZ treatment. **G, H** Heatmap showing quantifications of Olig2, Nkx 2.2, ASPA and CC1 in the GM (SSCx, MCx, Acc) and WM (Cc, Ec, Cg) at Bregma 0.8 and - 1.3 regions. The overall statistically significant differences across Bregma regions between control and cuprizone in each genotype is labelled by asterisks (WT black*; APP/PS1 red*) and the significant differences between WT and APP/PS1 is labelled by hash (WT black#; APP/PS1 red#) on right (n = 4 – 7, *p < 0.05; **p < 0.01; ^#^ p < 0.05; ^##^ p < 0.01; ^###^ p < 0.001; ^####^ p < 0.001). **I** The histology data indicating the higher vulnerability of mOL to CPZ in APP/PS1 was confirmed by gene expression analysis of whole brain tissue lysate. A distinct transcriptional response of OL (*Olig2*), OPC (*Sox10*, *Pdgfra*), mOL (*Cnpase*, *Myrf*, *Plp1*, *Mag*) and axon (tubb3) was detected where a significantly reduced *Cnpase* in APP/PS1 confirmed the higher susceptibility of mOL **J, K** Myelin proteins and OL cell density across the cortex and corpus callosum were plotted against the MTR and DTI measurements (FA, MD, AD and RD) and presented as heatmap. A distinct correlation pattern between WT and APP/PS1 animals was found, where most statistically significant correlations were found in corpus callosum. Significant correlation between PLP measurements against diffusivity was only found in the WT. Correlation analyzed by Pearson test. **L** MTR signal was highly correlated with the ASPA^+^ mOL density in the cortex and corpus callosum of APP/PS1 but only in the latter of WT. Two-way ANOVA followed by Tukey’s multiple comparisons test (n = 4 – 6, *p < 0.05; **p < 0.01, ***p < 0.001, ****p < 0.0001). Correlation analyzed by Pearson test with r and P values denoted on graph.

Gene expression analysis of the whole tissue lysate at Bregma -1.3 largely confirmed these immunohistochemical findings. As we used the entire brain for the analysis, the GM vs WM differences were lost. Nonetheless, we found that cuprizone increased *Olig2* and *Sox10* mRNA in WT, but not in APP/PS1, while it induced a significant reduction of *Cnpase*, a mOL marker, in APP/PS1 but not in WT. Intriguingly, cuprizone significantly increased *Myrf*, the master myelin gene transcription factor in APP/PS1 even as the expression of mature OL genes, *Plp1* or *Mag*, declined (Fig. 10I). In untreated animals, there were no significant difference observed between WT and APP/PS1. Next, we asked if the microscopic findings of OL density are directly proportional to the macroscopic changes detected by neuroimaging. The Pearson test between myelin fibers density, OL density, DTI and MTS measurements showed that such correlation differs between WT and APP/PS1 animals as well as between GM and WM (Fig. 10J, K). In the WM, there was a significant correlation between ASPA^+^ mOL density and MTR (Fig. 10L). By contrast, in the GM, while the mutant continued to show a significant correlation, the effect was lost in the WT. Together, our results suggest that the APP_SWE_/PSEN1_dE9_ transgenes do not affect the endogenous OL population but render it more vulnerable to cuprizone.

### Genotype- and cuprizone-dependent DNA damage in glial cells

To investigate the high vulnerability of OL population in APP/PS1 cortex, the DEG between WT and APP/PS1 cortices was analyzed (Fig. 11A). Among the DEGs identified at baseline (131 genes) and after cuprizone (144 genes), 49 and 22 genes respectively were up- or down-regulated in APP/PS1, regardless of treatment (Fig. 11B). Gene ontology analysis showed that the common upregulated DEGs were enriched in cell cycle-related processes (*Cell Cycle Process* GO:0022402), while the downregulated DEGs were enriched in metabolism-related processes (*Small Molecule Biosynthetic Process* GO:0044283) (Fig. 11C). Although the expression did not change with cuprizone treatment, *Atm* and *Mif* are two highly significant DEGs that may contribute to the higher OL vulnerability in APP/PS1 (Fig. 11D). Cuprizone-mediated copper chelation induces OL cell death by ferroptosis (79), which is the likely consequence of DNA damage (80, 81). Myelinating OLs are highly liable to DNA double strand break (DSB) repair (9, 21). *Atm* is the major DSB repair kinase and the significantly elevated *Atm* in APP/PS1 may reflect a response to more profound DNA damage in APP/PS1 OL after cuprizone treatment. *Mif (*macrophage migration inhibitory factor) is a pleiotropic cytokine that regulates neuroinflammation in multiple sclerosis (82, 83). Cuprizone significantly increased *Mif* in WT as expected, but the significantly lowered *Mif* level in APP/PS1 at baseline and after cuprizone treatment, suggests that the neuroinflammation in APP/PS1 is atypical. The important implication as we found 19 microglia-enriched or microglia associated genes from the targeted RNA-seq that differed between WT and APP/PS1 at these young ages, well before any diffuse amyloid plaque is formed (Fig. 11E).

**Figure 11.**
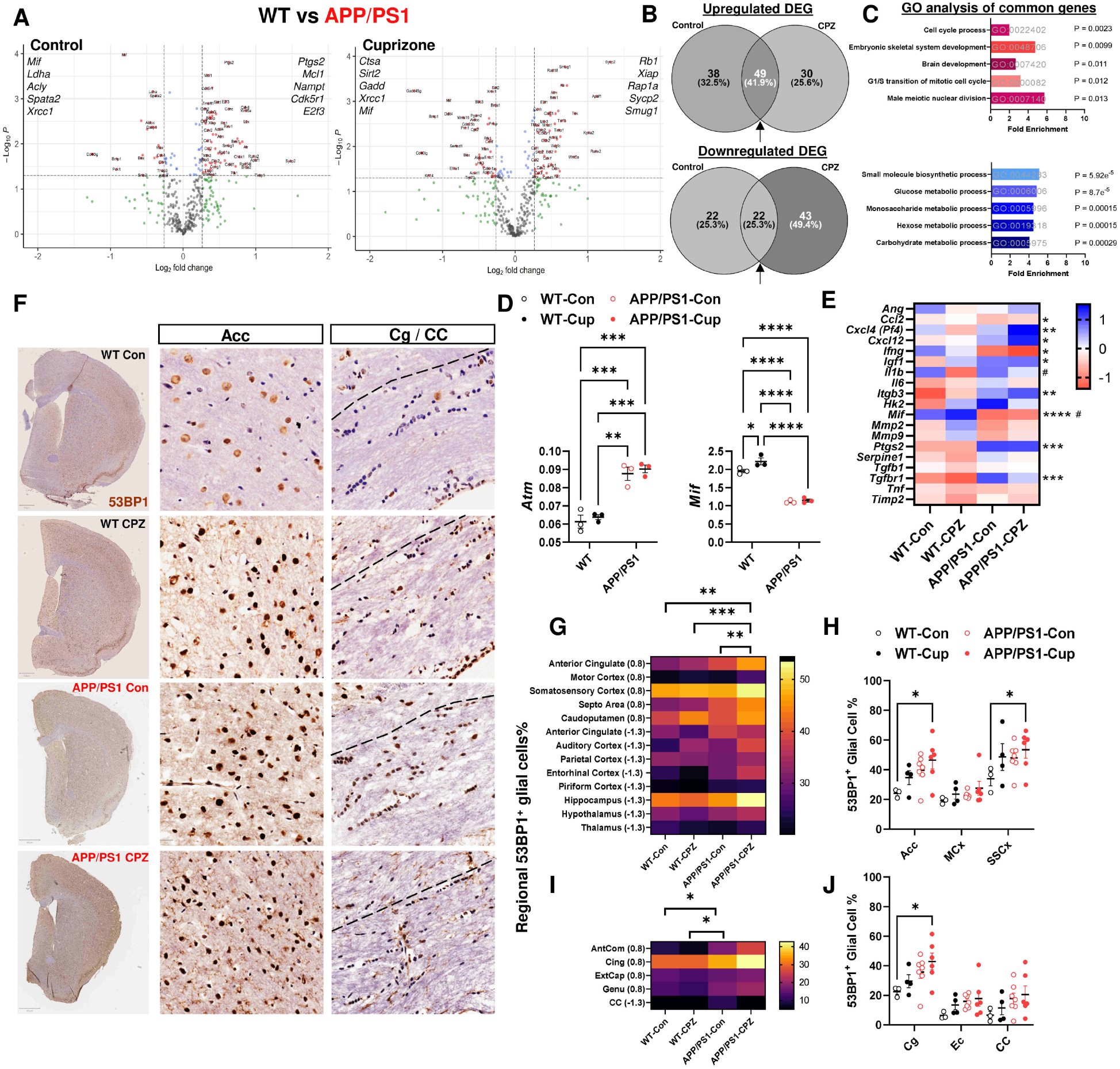
Robust DNA double strand breaks formation in APP/PS1 upon cuprizone-induced demyelination. **A** Volcano plot of targeted RNAseq of the cerebral cortices between WT and APP/PS1 showing the DEGs in control and cuprizone with the top five upregulated and downregulated genes stated. **B** Comparison between the DEGs in A identified 49 and 22 genes commonly upregulated and downregulated in APP/PS1 with or without cuprizone treatment. **C** Gene ontology analysis with PANTHER-based enrichment showing the top five biological processes significantly enriched in the DEGs commonly upregulated or downregulated in APP/PS1 vs WT. The GO term references and the enrichment significance are denoted on the graph. **D** Representative transcripts, *Atm* and *Mif*, from *Cell cycle process* (GO:0022402) and *Small molecule biosynthetic process* (GO:0044283) are shown, where both genes are differentially expressed in WT and APP/PS1. Of note, *Mif* is a cytokine that is important for demyelination-remyelination process, and it was significantly increased by CPZ in WT but not in APP/PS1. **E** All transcripts highly enriched in microglia/macrophage in the panel (19/416 genes) were plotted as heatmap. The difference between WT vs APP/PS1 (*) and between Con vs CPZ (#) are denoted and showed that the microglial response in APP/PS1 cortex is different from WT (*p < 0.05; **p < 0.01, ***p < 0.001, ****p < 0.0001; ^#^ p < 0.05). **F** Representative whole slide image of 53BP1 immunoreactivity(-IR) on coronal brain section at Bregma 0.8 in WT, R1.40 and APP/PS1 with or without CPZ treatment (*left*). The microscopic image of 53BP1-IR across Acc (middle) and Cg/CC (right) were shown. CPZ-induced a significant increase of 53BP1^+^ nucleus among neurons and glial cells in Acc and Cg/CC, respectively. Neuronal cells appeared as large nucleus with intense 53BP1-IR upon CPZ or in APP/PS1. Glial cells in Cg/CC appeared as a thread or bundle of small nucleus highly resemble of OLs. **G** Quantification across thirteen GM regions at Bregma 0.8 and -1.3 also showed a genotype- and CPZ-dependent increase of glial cell-like 53BP1^+^ nucleus. **H** The strongest increase of 53BP1^+^ glial cells was found in the Acc and SSCx, but no corresponding reduction of cell density was identified. **I, J** Quantification across five WM regions at Bregma 0.8 and -1.3 showed a genotype- and CPZ-dependent increase of 53BP1^+^ nucleus in these myelinated regions with the strongest effect found in the Cg. No overall cell density change was found in association with the 53BP1 signal. Statistical analysis between genotype or treatment in all regions performed by Two-way ANOVA repeated measures followed by Tukey’s multiple comparisons test; region specific changes was tested by Two-way ANOVA with Tukey’s multiple comparisons (* p < 0.05; **p < 0.01, ***p < 0.001, ****p < 0.0001).

The dysregulated DNA repair and neuroinflammation may therefore contribute to the high vulnerability of cortical OL population in APP/PS1. We immunolabelled DSBs with an antibody against 53BP1, a DNA damage associated protein that is a downstream substrate of ATM signaling and found a significant and global increase of 53BP1^+^ DSBs in glial cells in a genotype- and treatment-dependent fashion across GM and WM (Fig. 11F, G). In anterior cingulate cortex and somatosensory cortex, where a significant reduction of mOL was observed, cuprizone induced the formation of 53BP1^+^ DSBs in glial cells by 46.5% and 53.4%, respectively (Fig. 11H). In the WM, genotype-dependent (p < 0.05) and treatment-dependent global increases of 53BP1^+^ glial cells were similarly identified (Fig. 11I, J). As exemplified in the cingulum/corpus callosum, cells in thread-like clusters resembling typical OLs in WM were strongly 53BP1 positive after cuprizone treatment in both WT and APP/PS1 (Fig. 6F). Of note, nuclear 53BP1^+^ signals were readily detected in neuronal cells of WT at baselin. While the intensity of neuronal 53BP1^+^ was increased by cuprizone, no statistical significance was found. Unlike glial cells, DSBs are endogenously present in neurons in response to activity (84, 85) and a large variation was found among GM regions. The genotype and treatment associated increase of 53BP1^+^ glial cells agreed with RNA-seq results and suggested a perturbed DNA repair response in APP/PS1 animals that may contribute to the higher vulnerability of OL to demyelination (21).

## DISCUSSION

This study was undertaken to clarify the causal relationship between myelin pathology and the progression of sporadic AD. We report MRI findings showing that macroscopic myelin abnormalities are not a response to an increasing severity of amyloidosis in either R1.40 or APP/PS1 transgenic AD mouse models. Quite the contrary, we find that myelin loss precedes and may be the cause, rather than the consequence, of the progression of AD pathology.

Results from three independent mouse models are consistent in showing that the OL population is highly vulnerable to amyloid peptides (7, 8). Indeed, mouse AD models carrying an APP transgene either alone (R1,40; APP_SWE_ (9)), or in conjunction with a mutant *γ*-secretase (APPPS1; APP_SWE_ and PS1_L166P_ (10)) and a mutant human tau transgene (3xTg, APP_SWE_ PS1_M146V_ and Tau_P301L_ (11–13)) all exhibited myelination and/or OL degeneration. In this study, the R1.40 and APP/PS1 brains have similar expression of human APP_SWE_ but differ in the presence of PSEN1. The presence of the PSEN1 gene in APP/PS1 accelerates the appearance of amyloid deposits by many months (39), and according to the amyloid cascade hypothesis, would be expected to results in an earlier and a more profound demyelination. Instead, myelin tract integrity was more compromised in R1.40 animals even at early ages. First, these data suggest that it is the presence of excess APP expression, not the mutant PSEN1, that is the most likely driver of myelin abnormalities. Second, as the addition of PSEN1 does increase APP cleavage, these data suggest that increased amyloidosis is not the major contributor to myelin abnormalities, in agreement with earlier human studies (9).

Following up on this, we show that a four-week cuprizone-mediated acute demyelination regime causes a significant accumulation of intraneuronal amyloid in APP/PS1, but not in R1.40. Acute demyelination-induced axonal injury is known to disrupt the axonal transport of APP, and cause APP accumulation in the axonal spheroids (31, 86–89). The intraneuronal amyloid deposition in cuprizone-treated APP/PS1 is likely driven by PSEN1_dE9_ (Fig. 12), as in the R1.40 the absence of enhanced PSEN1 activity is correlated with the absence of significant amyloid pathology after demyelination. The accumulation of intraneuronal amyloid is an early neuropathology in AD and aging. In human AD, the earliest buildup of intraneuronal amyloid is found in neurons of the basal forebrain and locus coeruleus before any diffuse plaque is formed (40, 41). This abnormal buildup not only triggers neuroinflammation, but also interferes with neuronal excitability and dendritic spine maintenance (41, 90, 91). We suspect that these inflammatory and synaptic features may make independent contributions to the etiology of AD, thus underscoring the complexity of the dementia phenotype. Intraneuronal amyloid deposition is reported in 5xFAD model as early as 1.5 months of age (92, 93) and in APP/PS1 model at 4 months (41, 91). In both models, this intraneuronal amyloid deposition precedes the earliest appearance of the first diffuse plaque (2 and 6 months, respectively). The significantly stronger formation of intraneuronal amyloid in the frontal cortex upon demyelination, but not in hippocampus or posterior forebrain, tracks with the spatiotemporal progression of amyloid plaque formation in APP/PS1 at later stage (78).

**Figure 12.**
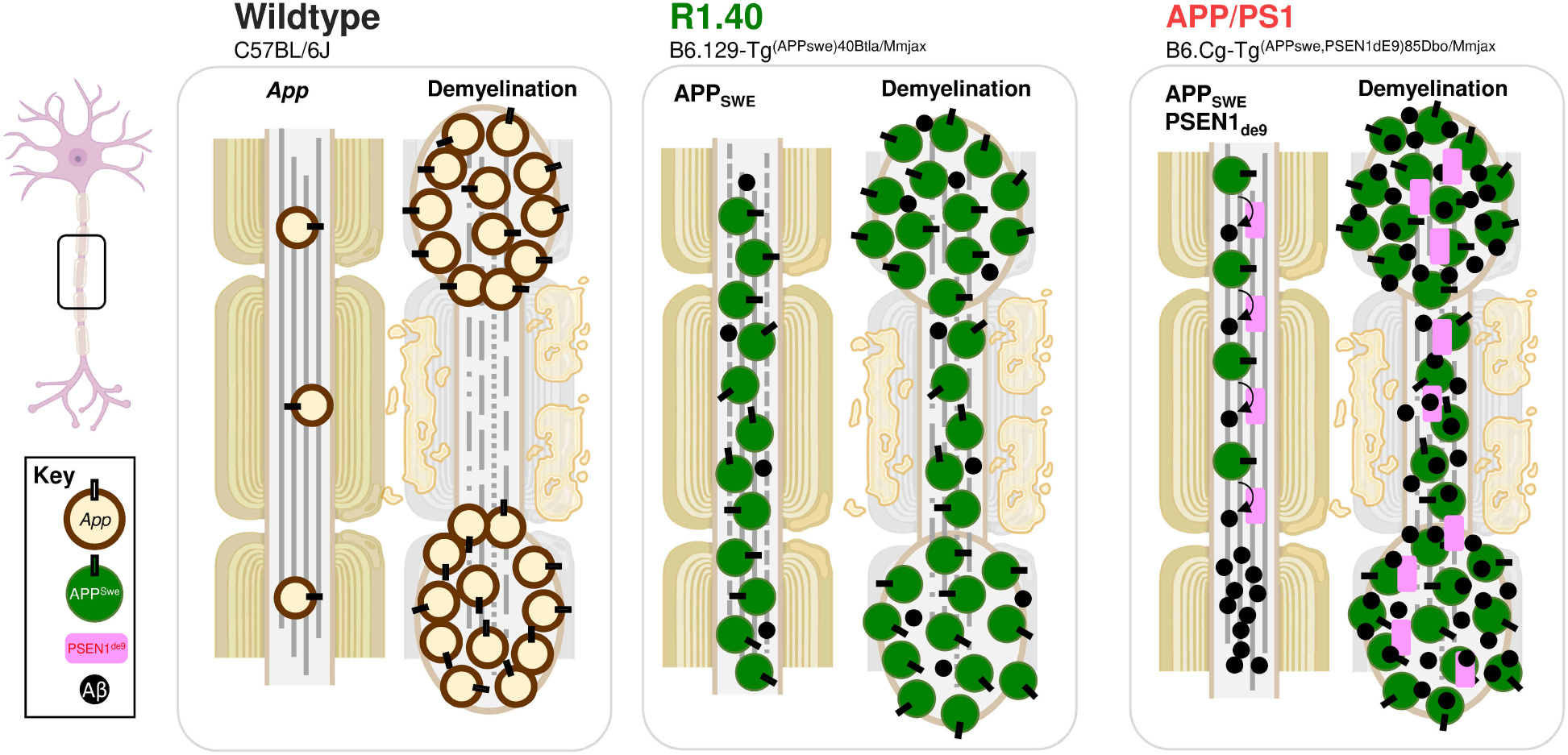
Putative cellular mechanism of demyelination driven amyloid accumulation in APP/PS1. The distribution of APP mediated by axonal transport, which is differentially impacted in WT, R1.40 and APP/PS1 animals in normal condition and after demyelination. In WT mice, App is distributed normally but demyelination caused the disruption of axonal transport and resulted in the accumulation of App spheroids (*App*, brown circles) along axons (26, 27). In R1.40 mice, as reported in similar models (69, 70), APPSWE overexpression (green circles) mice overloaded the axonal transport system even at the baseline level. Demyelination amplified such axonal deficit and the accumulation of APPSWE resulted in axonopathy with distinct DTI and MRS signatures in neuroimaging without any effects on amyloidosis. In APP/PS1 mice, although APPSWE expression is even higher than that in R1.40, the additional PSEN1dE9 mutant transgene (pink rectangle) accelerated the cleavage of APPSWE into Aβ (small black circles), and at least partially reversed the axonal overloading burden of APPSWE. It is plausible that demyelination in APPSWE, however, caused the accumulation of APPSWE and PSEN1dE9 at the axonal spheroids or the neuronal soma which served as the scaffold for amyloidosis. As a consequence, a significant level of amyloidosis occurred and resulted in an abnormal buildup of intraneuronal amyloid deposition (amyloidopathy) within four weeks of CPZ treatment. Future experiments are warranted to determine if demyelination will induce diffuse amyloid plaque formation and widespread neuronal injury after prolonged demyelination in R1.40 and APP/PS1.

The shifts in brain chemistry detected by MRS expanded this perspective considerably. Despite the absence of *PSEN1* gene, R1.40 animals showed the lowest level of NAA/Cr and Lipid/Cr (CH_2_ and CH_3_), both of which are clinically recognized markers for diffuse axonal damage in cerebral WM after traumatic brain injury in man and rodent (74, 94), while reduced Glx/Cr level and NAA/Cr are used as biomarkers of cortical demyelination in primary progressive MS patients (95). The suggestion is that R1.40 mice are highly vulnerable to both demyelination and axonal injury, compared with APP/PS1. In this regard it is worth recalling another feature of the APP transgene in R1.40. Unlike the APP/PS1 transgenes, which are cDNAs driven by neuron specific *Prnp* promoter, the R1.40 transgene is the entire human APP genomic sequence with all introns plus tens of kilobases of the 3’ and 5’ sequences inserted into the mouse genome on a yeast artificial chromosome (35). Thus, in R1.40 mice, APP will be expressed in all tissue and cell types, including OLs in a temporal progression that more closely mimics that found in human.

This concept is supported from our RNA-seq findings where R1.40 differs from APP/PS1 in genes responsible for metabolic processes, such as significantly upregulated *Acly* that is required to fuel augmented axonal transport with high metabolic demands (76). Our βIII-tubulin immunohistochemistry did not demonstrate the presence of acute axonal injury in any AD models at these relatively young ages. At later stages, axonopathy and defects in axon initial segments are found in APP_SWE_ overexpression models including R1.40 (77, 96). The axonal toxicity of APP overexpression has been attributed to its perturbation on axonal transport because overloaded APP may interrupt kinesin motor via JIP-1 (71). Indeed, human APP overexpression alone in mouse models even without familial AD related mutation (I5 vs J20 strains), contributes to AD-like phenotype including epileptic and synaptic loss independent of amyloidosis or amyloid plaque formation (72, 97, 98). In agreement with reports by Stokin et al (69, 70), APP/PS1 is spared from major axon transport defects likely due to the PSEN1_de9_ mutation. The amyloidosis is augmented by the PSEN1_dE9_ transgene in APP/PS1(43), but the accelerated cleavage of large APP protein into small Aβ peptide is also likely to ease axonal transport overloading (69). The distinct patterns of MRS and diffusivity in R1.40 after demyelination are therefore likely to be the consequence of APP overloading and subsequent myelin loss.

Our study is in line with a recent publication from Depp et al (99), in which diffuse amyloid plaque formation is reported in the cortex of 5xFAD and APP^NFGL^ mice after conditional genetic deletion of mOL or after cuprizone (4 weeks and 4 weeks of recovery). Consistent with their findings (99), we detected no alternation in the levels of the full-length APP protein level despite the significant increase of intraneuronal amyloid deposition in the cortex of APP/PS1. The suggestion is that it is the distribution of amyloid, not expression level of APP, that is disrupted by demyelination. In terms of behavioral changes, our findings also agree with their Y-Maze experiments (99). Both groups report no changes in spontaneous alternation were found despite increased activity. In the aggregate, the data suggest that acute demyelination accelerates amyloid pathology without imminent effects on cognitive functions.

Cortical demyelination can not only cause acute axonal injury in human MS and its experimental models (29, 89), but can also trigger the reorganization of cortical networks (100, 101). It is conceivable that such an adaptive response is recapitulated in cuprizone-treated R1.40 and resulting in axonal injury and cortical reorganization that leads to the unexpected MRI findings. Thus, interpretation of the biophysical underpinning of the unconventional DTI finding is speculative and warrants further experimental investigation (102).

Myelin loss in the frontal lobe is an early structural change in the aging human brain (103). The significant formation of intraneuronal amyloid deposition in the frontal cortex prompted us to ask if cortical OLs are more vulnerable in APP/PS1. In human AD brain, cortical OL degeneration is tightly associated with DSB formation independent of amyloid deposition or Braak stage (9, 18), and this OL-specific DNA damage is recapitulated in our cuprizone model. Both our RNA-seq and histology findings show that the density of *Atm* and 53BP-positive glial cells is significantly higher in the APP/PS1 cortex. This increased DNA damage response would be predicted since the OL lineage is highly sensitive to DNA damage (21). Coupled with the reported high turnover rate of myelin in the APP/PS1 strain (104), it is likely the increased level of DNA damage leads to cell cycle-associated OL cell death (20, 105). The enhanced expression of cell cycle-related genes and the significant increase of OL/OPC-enriched genes *Rap1a*, *Bnip3l*, *Desi1* and *Tmeff1* in APP/PS1 cortex support this argument, but the cellular mechanism remains elusive.

Biophysically, the detection of myelin changes by traditional MRI-neuroimaging is based on the physical property of atomic nuclei in a strong magnetic field. The mathematical calculation of the magnetization of water, protons or macromolecules are reflected as a measurement of the fluid diffusion or chemical species in brain structures and serve as surrogates of myelin tract integrity (106). Biologically, the internodal myelin segment between nodes of Ranvier can be produced and maintained by a pre-existing mOL, and/or a mOL newly derived from an OPC (107–109). The cellular dynamics between the existing myelinating mOL and their eventual replacement by differentiating OPC leads to the variable myelin thickness among brain regions detectable by DTI (110, 111). Earlier reports correlated histological loss of myelin tracts with DTI (64, 65, 112–114), MTR (66, 115, 116) and MRS (117–119) based myelin protein immunohistochemistry, myelin-specific stain or transmission electron microscopy. These snapshots of radiopathological correlation in limited regions cannot capture the dynamics of OL differentiation. (120). Furthermore, myelin proteins, such as PLP and MOG, differs substantially even in the same region and it complicates the interpretation of earlier studies (121–123). Here, our results showed that MTR, but not DTI is more robustly associated with OL dynamics, and this correlation differs between APP/PS1 and WT. Given the strong correlation between intraneuronal amyloid formation and mean diffusivity in the cortex, cautious DTI interpretation is required at least for experimental models. Indeed, the demise of OL may result in a substantial re-organization of the myeloarchitecture without affecting the overall myelin content, further complicating DTI data interpretation (9, 18, 124, 125). MRS may provide a closer account of myelin content by revealing chemical shifts, such as lipids, in MS patients (126–130). By using CEST (Chemical exchange saturation transfer)-MRI, future study may capture the OL dynamics more accurately by targeting specific myelin lipid or amide shift (131–133).

## CONCLUSION

Overall, the data in this study show that the character of cuprizone-mediated demyelination is highly dependent on genetic background. Wildtype, APP_SWE_ and APP_SWE_/PSEN1_dE9_ transgenic mice differ substantially in their MRI signature, their myelin tract integrity and their levels of intraneuronal amyloid deposition. These divergent responses lead us to propose a model of Alzheimer’s disease in which myelin degradation, beginning with that found during normal aging, could be a cause rather than a consequence of pathological progression of the dementia.

## DECLARATIONS

### Ethics approval and consent to participate

- This study does not involve human data or tissue
- This study involves animals

`o` All animal experiments in this study are approved by Animal Subjects Ethics Sub-committee (ASESC), Research Committee, The Hong Kong Polytechnic University with ASESC reference number 19-20/6-HTI-R-HMRF

### Consent for publication

- Not applicable

### Availability of data and material

- The datasets during and/or analyzed during the current study available from the corresponding author on reasonable request.

### Acknowledgments

- The present work is generously supported by Health and Medical Research Fund from Food and Health Bureau (HMRF05163736) and Early Career Scheme from Research Grant Council (ECS25104220), Hong Kong Special Administrative Region. Dr. Kai-Hei Tse is also supported by PolyU Start-up Fund P0030307. Prof. Karl Herrup is supported by Start-up Fund at Department of Neurobiology, University of Pittsburgh and the Australian National Health and Medical Research Fund (APP1160691), the Pennsylvania Department of State (4100087331) and additional support from the NIA (R01 AG069912).
- The authors would like to acknowledge medical technologists, Manfred Hon-Man Lai, Rick Wing-Ho Ma and Laura Chi-Fan Yuen at the Tai Tung Veterinary Laboratory, Agriculture, Fisheries and Conservation Department of HKSAR government for their kind technical support in digital pathology scanning.

### Authors’ contributions

- G.W.Y.C, I.W.T.M, and J.H performed animal experiments including neuroimaging, histopathology, molecular tests with data acquisition and analysis with equal contribution. S.H.S.Y performed the molecular test. P.H performed the histopathology test. Z.C performed the neuroimaging data acquisition. K.H.T wrote the manuscript, supervised the study and obtained funding. H.K.H.M, K.H, K.W.Y.C and K.H.T conceptualized the study, designed the experiment, and edited the manuscript. All authors read, critically reviewed, and approved the manuscript.

## ACKNOWLEDGEMENTS

The present work is generously supported by Health and Medical Research Fund from Food and Health Bureau (HMRF05163736) Hong Kong Special Administrative Region. Dr. Kai-Hei Tse is also supported by PolyU Start-Up Fund (P0030307) and Early Career Scheme from Research Grant Council (ECS25104220). Prof. Karl Herrup is supported by a start-up fund at Department of Neurobiology, University of Pittsburgh, the Australian National Health and Medical Research Fund (APP1160691) with additional support from the NIA (R01 AG069912) and NINDS (NS120922).

