## Supplementary material for "Cuprizone drives divergent neuropathological changes in different mouse models of Alzheimer’s disease": Table S1

### Table S1 – Differential Expression Gene Lists of targeted RNAseq

| Supplementary Table S1A – Differential Gene Expression of WT-Con vs WT-CPZ | | | | | |
|---|---|---|---|---|---|
| **Symbol** | **Fold Regulation** | **adj. p-Value** | **Symbol** | **Fold Regulation** | **adj. p-Value** |
| *Ccnd2* | 1.17 | 0.002208 | *Ctgf* | 0.86 | 0.008216 |
| *Fas* | 1.22 | 0.004535 | *Nrp1* | 0.86 | 0.009746 |
| *Hk3* | 1.52 | 0.004639 | *Casp1* | 0.67 | 0.011278 |
| *Cdkn3* | 1.3 | 0.005656 | *Rad18* | 0.88 | 0.014898 |
| *Cdkn1a* | 1.76 | 0.007162 | *Rif1* | 0.76 | 0.018372 |
| *Brca1* | 1.34 | 0.007616 | *Il1b* | 0.6 | 0.027517 |
| *Fn1* | 1.4 | 0.007957 | *Pot1a* | 0.86 | 0.033357 |
| *Txnip* | 1.8 | 0.00825 | *Efnb2* | 0.83 | 0.035459 |
| *Gadd45a* | 1.43 | 0.009395 | *Figf* | 0.81 | 0.036166 |
| *Plk1* | 1.98 | 0.009502 | *Mmp14* | 0.84 | 0.03981 |
| *Ppp1r15a* | 1.21 | 0.01073 | *Ercc2* | 0.89 | 0.041664 |
| *Gadd45g* | 1.77 | 0.012235 | | | |
| *Egf* | 2 | 0.023304 | | | |
| *Bbc3* | 1.35 | 0.031022 | | | |
| *Pfkfb3* | 1.23 | 0.031305 | | | |
| *Mcl1* | 1.08 | 0.034944 | | | |
| *Casp7* | 1.42 | 0.037105 | | | |
| *Anxa2* | 1.18 | 0.049072 | | | |
| *Pdk4* | 1.4 | 0.049128 | | | |

| Supplementary Table S1B – Differential Gene Expression of R140-Con vs R140-CPZ | | | | | |
|---|---|---|---|---|---|
| **Symbol** | **Fold Regulation** | **adj. p-Value** | **Symbol** | **Fold Change** | **adj. p-Value** |
| *Mki67* | 1.79 | 0.002795 | *Terf2* | 0.97 | 0.000472 |
| *Brca2* | 1.35 | 0.007449 | *Casp3* | 0.85 | 0.000594 |
| *Msn* | 1.34 | 0.007578 | *Snai2* | 0.72 | 0.006256 |
| *Fas* | 1.3 | 0.007788 | *Sirt1* | 0.77 | 0.006789 |
| *Cdc20* | 1.35 | 0.015065 | *Xrcc3* | 0.77 | 0.017994 |
| *Rad50* | 1.22 | 0.021573 | *Fgf1* | 0.87 | 0.018695 |
| *Gpd2* | 1.15 | 0.025295 | *Tmeff1* | 0.81 | 0.022047 |
| *Neil3* | 2.15 | 0.025533 | *Rad23a* | 0.88 | 0.029049 |
| *Edn1* | 1.54 | 0.026942 | *Itgb3* | 0.69 | 0.039115 |
| *Slc2a1* | 1.44 | 0.032357 | *Neil1* | 0.77 | 0.04025 |
| *Cdh1* | 1.35 | 0.033395 | *Ndrg1* | 0.85 | 0.043209 |
| *Casp6* | 1.2 | 0.036142 | *Apaf1* | 0.71 | 0.043725 |
| *Ahnak* | 1.25 | 0.037464 | *Rad1* | 0.9 | 0.044418 |
| *Anxa2* | 1.34 | 0.037993 | *Fbrsl1* | 0.84 | 0.045113 |
| *Col18a1* | 1.55 | 0.040086 | *Bhlhe40* | 0.75 | 0.046802 |
| *Brip1* | 1.64 | 0.040182 | | | |
| *Mgmt* | 1.09 | 0.047893 | | | |
| *Rnf8* | 1.19 | 0.048021 | | | |

| Supplementary Table S1C – Differential Gene Expression of APPPS1-Con_V_APPPS1-CPZ | | | | | |
|---|---|---|---|---|---|
| **Symbol** | **Fold Change** | **adj. p-Value** | **Symbol** | **Fold Change** | **adj. p-Value** |
| *Birc3* | 1.17 | 0.005564 | *Uqcrfs1* | 0.93 | 0.002307 |
| *Pf4* | 1.26 | 0.008069 | *Sirt2* | 0.87 | 0.00342 |
| *Aurkb* | 1.43 | 0.01178 | *Ctgf* | 0.8 | 0.004707 |
| *Efnb2* | 1.1 | 0.014194 | *Car9* | 0.48 | 0.012089 |
| *Ercc2* | 1.09 | 0.014641 | *Ccnc* | 0.91 | 0.015818 |
| *Ccnt1* | 1.14 | 0.016326 | *Slc2a1* | 0.91 | 0.016676 |
| *Neil1* | 1.13 | 0.017654 | *Pim1* | 0.78 | 0.021984 |
| *Mad2l1* | 1.18 | 0.01801 | *Tmeff1* | 0.87 | 0.038836 |
| *Dll4* | 1.65 | 0.01832 | *Ndrg1* | 0.79 | 0.040795 |
| *Vegfc* | 1.12 | 0.029534 | | | |
| *Cdh5* | 1.18 | 0.035733 | | | |
| *Casp6* | 1.14 | 0.039957 | | | |
| *Tnfrsf11b* | 1.46 | 0.040566 | | | |
| *Epo* | 2.9 | 0.043024 | | | |
| *Txnip* | 1.22 | 0.043793 | | | |
| *Mmp9* | 1.18 | 0.045605 | | | |
| *Adora2b* | 1.18 | 0.048271 | | | |
| *Kdr* | 1.35 | 0.050004 | | | |

Table S1 – Differential Expression Gene Lists of targeted RNAseq

| **Supplementary Table S1D – Differential Gene Expression of WT-Con vs R140-Con** | | | | | |
|---|---|---|---|---|---|
| **Symbol** | **Fold Change** | **adj. p-Value** | **Symbol** | **Fold Change** | **adj. p-Value** |
| *Xpc* | 1.41 | 0.000328 | *Mif* | 0.68 | 0.000515 |
| *Pnkp* | 1.2 | 0.001051 | *Gpd2* | 0.79 | 0.000605 |
| *E2f3* | 1.19 | 0.001331 | *Nop10* | 0.61 | 0.001334 |
| *Egf* | 2.34 | 0.002092 | *Tspan13* | 0.62 | 0.002002 |
| *Mtor* | 1.17 | 0.004179 | *Fam162a* | 0.68 | 0.005439 |
| *Neil1* | 1.47 | 0.004781 | *Cks1b* | 0.75 | 0.006147 |
| *Casp3* | 1.35 | 0.00491 | *Pgk1* | 0.86 | 0.007533 |
| *Bmi1* | 1.21 | 0.00499 | *Snai1* | 0.66 | 0.007581 |
| *Cdk5r1* | 1.26 | 0.009381 | *Bccip* | 0.89 | 0.012284 |
| *Kntc1* | 1.54 | 0.009477 | *Pinx1* | 0.74 | 0.015109 |
| *Zbtb22* | 1.15 | 0.010145 | *Aldoa* | 0.88 | 0.015424 |
| *Ddb2* | 1.46 | 0.011847 | *Mxi1* | 0.8 | 0.015551 |
| *Itgb3* | 1.83 | 0.01354 | *Lgals3* | 0.37 | 0.015749 |
| *Akt1* | 1.06 | 0.015366 | *Cdk5rap1* | 0.92 | 0.017797 |
| *Cdc25a* | 1.31 | 0.02192 | *Ing1* | 0.82 | 0.020241 |
| *Rad23a* | 1.2 | 0.02235 | *Rgs2* | 0.82 | 0.022624 |
| *Sirt1* | 1.3 | 0.022609 | *Cib1* | 0.84 | 0.023118 |
| *Map2k1* | 1.13 | 0.024702 | *Cks2* | 0.81 | 0.024027 |
| *Acly* | 1.13 | 0.024715 | *Cdk8* | 0.88 | 0.02882 |
| *Eno2* | 1.22 | 0.024795 | *Polb* | 0.88 | 0.033407 |
| *Fbrsl1* | 1.41 | 0.02866 | *Angpt1* | 0.75 | 0.033604 |
| *Pcx* | 1.15 | 0.03243 | *Ldha* | 0.93 | 0.035295 |
| *Cdh2* | 1.17 | 0.033051 | *Rnf8* | 0.9 | 0.036124 |
| *Ccnd3* | 1.21 | 0.033607 | *Ppil2* | 0.88 | 0.036795 |
| *Slc2a3* | 1.17 | 0.033817 | *Rpa3* | 0.74 | 0.038942 |
| *Casp2* | 1.33 | 0.035984 | *Cyld* | 0.87 | 0.044188 |
| *Timp2* | 1.05 | 0.036359 | *Ang* | 0.77 | 0.045973 |
| *Wnt5a* | 1.78 | 0.037132 | | | |
| *Hk1* | 1.05 | 0.037279 | | | |
| *Serpinf1* | 1.18 | 0.037697 | | | |
| *Tnks2* | 1.13 | 0.038032 | | | |
| *Xiap* | 1.12 | 0.039263 | | | |
| *Wee1* | 1.12 | 0.039489 | | | |
| *Cdkn2d* | 1.09 | 0.042037 | | | |
| *Rev1* | 1.2 | 0.046641 | | | |
| *Tgfbr1* | 1.25 | 0.04697 | | | |
| *Igfbp5* | 1.3 | 0.048914 | | | |

### Table S1 – Differential Expression Gene Lists of targeted RNAseq

**Supplementary Table S1E – Differential Gene Expression of WT-Con vs APP-Con**

| Symbol | Fold Change | adj. p-Value | Symbol | Fold Change | adj. p-Value |
|---|---|---|---|---|---|
| *Ptgs2* | 1.55 | 0.000156 | *Mif* | 0.57 | 0.000105 |
| *Mcl1* | 1.27 | 0.000302 | *Ldha* | 0.75 | 0.000695 |
| *Nampt* | 1.32 | 0.000569 | *Acly* | 0.88 | 0.000725 |
| *Cdk5r1* | 1.26 | 0.000587 | *Spata2* | 0.77 | 0.000854 |
| *E2f3* | 1.5 | 0.001158 | *Xrcc1* | 0.67 | 0.003057 |
| *Sirt1* | 1.39 | 0.001183 | *Aldoc* | 0.73 | 0.00337 |
| *Tmeff1* | 1.27 | 0.001261 | *Xab2* | 0.75 | 0.00423 |
| *Cdk6* | 1.81 | 0.001548 | *Casp1* | 0.74 | 0.004267 |
| *Terf2ip* | 1.14 | 0.001642 | *Ccnt1* | 0.77 | 0.004295 |
| *Cdh2* | 1.23 | 0.001669 | *Ctsa* | 0.71 | 0.004715 |
| *Ccnc* | 1.55 | 0.001675 | *Pdk2* | 0.79 | 0.010778 |
| *Wnt5a* | 1.82 | 0.001824 | *Msn* | 0.8 | 0.012016 |
| *Bnip3l* | 1.31 | 0.001952 | *Ppil2* | 0.9 | 0.016529 |
| *Tgfbr1* | 1.33 | 0.001973 | *Cd40lg* | 0.42 | 0.016986 |
| *Uqcrfs1* | 1.29 | 0.002059 | *Epo* | 0.42 | 0.016986 |
| *Rad17* | 1.37 | 0.002167 | *Gsc* | 0.42 | 0.016986 |
| *Xpc* | 1.32 | 0.002403 | *Ifng* | 0.42 | 0.016986 |
| *Wee1* | 1.29 | 0.00251 | *Il1rn* | 0.42 | 0.016986 |
| *Bccip* | 1.3 | 0.002673 | *Mmp3* | 0.42 | 0.016986 |
| *Mnat1* | 1.3 | 0.002724 | *Pgk2* | 0.42 | 0.016986 |
| *Pms1* | 1.24 | 0.002954 | *Snai1* | 0.68 | 0.017722 |
| *Ctgf* | 1.22 | 0.003125 | *Cdk4* | 0.81 | 0.017765 |
| *Lox* | 1.83 | 0.003195 | *Pcx* | 0.81 | 0.017942 |
| *Xiap* | 1.32 | 0.003413 | *Zbtb22* | 0.92 | 0.018274 |
| *Rb1* | 1.48 | 0.003446 | *Pgam2* | 0.78 | 0.019267 |
| *Cul3* | 1.24 | 0.003541 | *Gpd2* | 0.89 | 0.020241 |
| *Igfbp4* | 1.13 | 0.003616 | *Mms19* | 0.76 | 0.020504 |
| *Rad1* | 1.17 | 0.003765 | *Bax* | 0.67 | 0.020572 |
| *Rev1* | 1.54 | 0.004083 | *Ercc2* | 0.85 | 0.020646 |
| *Acsl4* | 1.41 | 0.004697 | *Bmp1* | 0.53 | 0.021263 |
| *Cav2* | 1.27 | 0.004958 | *Cdk5rap1* | 0.85 | 0.021521 |
| *Rad23b* | 1.33 | 0.006176 | *Neil1* | 0.84 | 0.021636 |
| *Bcl2* | 1.37 | 0.00622 | *Cib1* | 0.82 | 0.025361 |
| *Atm* | 1.43 | 0.006831 | *Sparc* | 0.85 | 0.025515 |
| *Met* | 1.68 | 0.00764 | *Tinf2* | 0.8 | 0.026288 |
| *Cul1* | 1.26 | 0.008451 | *Dffa* | 0.85 | 0.027342 |
| *Bmi1* | 1.65 | 0.008839 | *Atp5a1* | 0.89 | 0.029575 |
| *Ccnd1* | 1.76 | 0.008931 | *Pkm* | 0.94 | 0.035639 |
| *Vcan* | 1.31 | 0.009121 | *Nbn* | 0.94 | 0.035751 |
| *Itgb3* | 1.72 | 0.009712 | *Pck1* | 0.53 | 0.036319 |
| *Smug1* | 1.54 | 0.01083 | *Thap3* | 0.82 | 0.038804 |
| *Atr* | 1.82 | 0.011645 | *Rad51* | 0.86 | 0.046166 |
| *Sp1* | 1.19 | 0.012251 | *Snai2* | 0.68 | 0.048822 |
| *Cul2* | 1.27 | 0.012556 | *Aldoa* | 0.83 | 0.049908 |
| *Skp2* | 1.36 | 0.012738 | | | |
| *Rad21* | 1.17 | 0.013304 | | | |
| *Ptges3* | 1.16 | 0.013512 | | | |
| *Gadd45a* | 1.29 | 0.013671 | | | |
| *Rap1a* | 1.48 | 0.015272 | | | |
| *Tfrc* | 1.4 | 0.016093 | | | |
| *Terf1* | 1.44 | 0.016696 | | | |
| *Mlh3* | 1.26 | 0.018046 | | | |
| *Runx2* | 1.96 | 0.018741 | | | |
| *Chek1* | 1.71 | 0.019184 | | | |
| *Rad23a* | 1.12 | 0.020626 | | | |
| *Casp2* | 1.2 | 0.021278 | | | |
| *Igfbp5* | 1.22 | 0.022292 | | | |
| *Ercc5* | 1.18 | 0.022438 | | | |
| *Apaf1* | 1.99 | 0.022788 | | | |
| *Vps13a* | 1.49 | 0.023459 | | | |
| *Cks2* | 1.36 | 0.023842 | | | |
| *Sycp2* | 2.79 | 0.023843 | | | |
| *Pms2* | 1.17 | 0.025962 | | | |
| *Fbrsl1* | 1.43 | 0.026095 | | | |
| *E2f1* | 1.29 | 0.026148 | | | |
| *Map3k2* | 1.36 | 0.026851 | | | |
| *Fgf2* | 1.31 | 0.026921 | | | |
| *Rad54l* | 1.81 | 0.027734 | | | |
| *Cald1* | 1.56 | 0.028409 | | | |
| *Tcf4* | 1.13 | 0.029323 | | | |
| *Rad18* | 1.24 | 0.029472 | | | |
| *Prkdc* | 1.65 | 0.031596 | | | |
| *Ccng1* | 1.36 | 0.031957 | | | |
| *Brca2* | 1.83 | 0.032752 | | | |
| *Slc16a3* | 1.3 | 0.035431 | | | |
| *Pfkfb4* | 1.16 | 0.035459 | | | |
| *Polb* | 1.18 | 0.038739 | | | |
| *Tfdp1* | 1.28 | 0.039064 | | | |
| *Mlh1* | 1.24 | 0.04181 | | | |
| *Hus1* | 1.21 | 0.042095 | | | |
| *Plk1* | 1.63 | 0.043929 | | | |
| *Bhlhe40* | 1.29 | 0.047245 | | | |
| *Blm* | 1.11 | 0.047512 | | | |
| *Tfdp2* | 1.13 | 0.048342 | | | |
| *Prkaa1* | 1.07 | 0.048431 | | | |
| *Timp1* | 2.08 | 0.049347 | | | |
| *Casp7* | 1.32 | 0.049499 | | | |

### Table S1 – Differential Expression Gene Lists of targeted RNAseq

| Supplementary Table S1F – Differential Gene Expression of R140-Con_vs_APP-Con | | | | | |
|---|---|---|---|---|---|
| **Symbol** | **Fold Change** | **adj. p-Value** | **Symbol** | **Fold Change** | **adj. p-Value** |
| *Bccip* | 1.43 | 6.07553E-06 | *Neil1* | 0.56 | 0.000298 |
| *Pgk1* | 1.26 | 0.000245828 | *Mtor* | 0.81 | 0.00056 |
| *Uqcrfs1* | 1.28 | 0.000478077 | *Ctsa* | 0.71 | 0.001396 |
| *Ccnc* | 1.45 | 0.000767603 | *Cpt2* | 0.78 | 0.001441 |
| *Polb* | 1.31 | 0.00089782 | *Mms19* | 0.71 | 0.003542 |
| *Cul1* | 1.23 | 0.00102114 | *Pcx* | 0.68 | 0.003659 |
| *Nampt* | 1.28 | 0.001514147 | *Acly* | 0.76 | 0.003902 |
| *Ptgs2* | 1.69 | 0.001586795 | *Slc2a4* | 0.61 | 0.004034 |
| *Tmeff1* | 1.2 | 0.001697291 | *Zbtb22* | 0.78 | 0.00446 |
| *Map3k2* | 1.38 | 0.002241199 | *Hk1* | 0.88 | 0.004606 |
| *Rap1a* | 1.69 | 0.002485362 | *Aldoc* | 0.68 | 0.005163 |
| *Rad23b* | 1.29 | 0.002489663 | *Map2k1* | 0.81 | 0.005597 |
| *E2f3* | 1.23 | 0.003797618 | *Ccnt1* | 0.72 | 0.005681 |
| *Tspan13* | 1.44 | 0.004151668 | *Trp73* | 0.71 | 0.005779 |
| *Bnip3l* | 1.33 | 0.004684623 | *Fgf1* | 0.78 | 0.005861 |
| *Ptges3* | 1.2 | 0.004742017 | *Ldha* | 0.8 | 0.006132 |
| *Cul2* | 1.25 | 0.0053982 | *Gadd45g* | 0.67 | 0.006455 |
| *Xiap* | 1.15 | 0.005403811 | *Mif* | 0.81 | 0.007065 |
| *Rev1* | 1.25 | 0.005438905 | *Parp3* | 0.65 | 0.009209 |
| *Mlh3* | 1.25 | 0.005982852 | *Xrcc1* | 0.73 | 0.009568 |
| *Cdk6* | 1.62 | 0.006888648 | *Spata2* | 0.67 | 0.00985 |
| *Pms1* | 1.31 | 0.006926681 | *Anapc2* | 0.86 | 0.010011 |
| *Vps13a* | 1.57 | 0.0070136 | *Cdc25a* | 0.8 | 0.010123 |
| *Nop10* | 1.39 | 0.007050302 | *Msn* | 0.77 | 0.01031 |
| *Ctgf* | 1.31 | 0.007298795 | *Epo* | 0.33 | 0.010513 |
| *Terf1* | 1.46 | 0.00811165 | *Pnkp* | 0.71 | 0.013406 |
| *Vcan* | 1.25 | 0.008553179 | *Dffa* | 0.81 | 0.013529 |
| *Cks2* | 1.64 | 0.008991708 | *Rad9a* | 0.73 | 0.016068 |
| *Atr* | 1.49 | 0.009245266 | *Xab2* | 0.66 | 0.016465 |
| *Rb1* | 1.34 | 0.010899322 | *Bmp1* | 0.52 | 0.018227 |
| *Cdk8* | 1.29 | 0.011071056 | *Pck1* | 0.42 | 0.018708 |
| *Fas* | 1.29 | 0.012602748 | *Vdac1* | 0.86 | 0.021028 |
| *Gpd2* | 1.09 | 0.013397222 | *Lig1* | 0.67 | 0.021184 |
| *Birc2* | 1.25 | 0.014157687 | *Xrcc3* | 0.75 | 0.022734 |
| *Mnat1* | 1.36 | 0.014364667 | *Casp3* | 0.82 | 0.022788 |
| *Desi1* | 1.18 | 0.015771089 | *Tinf2* | 0.83 | 0.022815 |
| *Rad21* | 1.12 | 0.016861946 | *Poll* | 0.81 | 0.024381 |
| *Met* | 1.46 | 0.017135677 | *Ercc2* | 0.75 | 0.024428 |
| *Rad50* | 1.49 | 0.017449375 | *Mad2l2* | 0.81 | 0.027706 |
| *Id1* | 1.49 | 0.018055356 | *Atp5a1* | 0.87 | 0.032785 |
| *Pinx1* | 1.34 | 0.018396587 | *Eno2* | 0.84 | 0.035323 |
| *Bmi1* | 1.33 | 0.019835476 | *Sirt6* | 0.71 | 0.035537 |
| *Cul3* | 1.17 | 0.021000769 | *Pdk2* | 0.86 | 0.035551 |
| *Hgf* | 1.42 | 0.021587305 | *Sertad1* | 0.79 | 0.038911 |
| *Rad17* | 1.36 | 0.023138992 | *Cdc25b* | 0.65 | 0.044615 |
| *Tfdp1* | 1.33 | 0.023242655 | *Gpi1* | 0.82 | 0.046648 |
| *Fam162a* | 1.29 | 0.02447702 | *Ddit4* | 0.63 | 0.047172 |
| *Cav2* | 1.25 | 0.02682842 | *Lig3* | 0.83 | 0.0488 |
| *Gadd45a* | 1.25 | 0.027342715 | | | |
| *Ccng1* | 1.36 | 0.027774061 | | | |
| *Rpa3* | 1.39 | 0.027984835 | | | |
| *Lgals3* | 2.85 | 0.029091767 | | | |
| *Acsl4* | 1.27 | 0.032335377 | | | |
| *Pms2* | 1.09 | 0.033330177 | | | |
| *Timp1* | 2.34 | 0.033366774 | | | |
| *Terf2ip* | 1.12 | 0.033798477 | | | |
| *Angpt1* | 1.48 | 0.03490973 | | | |
| *Sycp2* | 2.26 | 0.035370614 | | | |
| *Ccnh* | 1.26 | 0.03551375 | | | |
| *Bnip3* | 1.22 | 0.036231805 | | | |
| *Tcf4* | 1.13 | 0.03752238 | | | |
| *Ccnd2* | 1.21 | 0.038111642 | | | |
| *Brca2* | 1.76 | 0.03814838 | | | |
| *Ccnb2* | 1.3 | 0.041126091 | | | |
| *Mad2l1* | 1.26 | 0.046454565 | | | |

Table S1 – Differential Expression Gene Lists of targeted RNAseq

| Supplementary Table S1G – Differential Gene Expression of WT-CPZ vs R140-CPZ | | | | | |
|---|---|---|---|---|---|
| **Symbol** | **Fold Change** | **adj. p-Value** | **Symbol** | **Fold Change** | **adj. p-Value** |
| *Mad2l2* | 1.16 | 0.004312 | *Ccnd2* | 0.76 | 0.003295 |
| *Mcm5* | 1.19 | 0.00957 | *Gadd45a* | 0.67 | 0.005388 |
| *Slc2a3* | 1.44 | 0.014572 | *Pgam2* | 0.62 | 0.005575 |
| *Hk2* | 1.55 | 0.014694 | *Tcf4* | 0.83 | 0.007385 |
| *Tek* | 1.16 | 0.014764 | *Fas* | 0.91 | 0.007467 |
| *Cdk5r1* | 1.21 | 0.015414 | *Fam162a* | 0.67 | 0.011585 |
| *Cald1* | 1.16 | 0.018314 | *Polb* | 0.78 | 0.014874 |
| *Slc2a4* | 1.84 | 0.01965 | *Desi1* | 0.82 | 0.015277 |
| *Cdk2* | 1.26 | 0.020133 | *Rpa3* | 0.71 | 0.015665 |
| *Vdac1* | 1.16 | 0.022938 | *Mif* | 0.72 | 0.015719 |
| *Eno2* | 1.23 | 0.0294 | *G6pdx* | 0.89 | 0.016514 |
| *Kpna2* | 1.63 | 0.036832 | *Nop10* | 0.58 | 0.017925 |
| *Msn* | 1.16 | 0.038004 | *Gadd45g* | 0.62 | 0.019663 |
| *Cd40lg* | 1.27 | 0.038533 | *Cpt2* | 0.84 | 0.020801 |
| *G6pc* | 1.27 | 0.038533 | *Xrcc6* | 0.82 | 0.024702 |
| *Gsc* | 1.27 | 0.038533 | *Pfkfb3* | 0.8 | 0.024808 |
| *Ifng* | 1.27 | 0.038533 | *Sirt1* | 0.94 | 0.027377 |
| *Mmp3* | 1.27 | 0.038533 | *Cdkn3* | 0.75 | 0.028145 |
| *Pck1* | 1.27 | 0.038533 | *Tspan13* | 0.6 | 0.030589 |
| *Pgk2* | 1.27 | 0.038533 | *Mnat1* | 0.84 | 0.03082 |
| *Pklr* | 1.27 | 0.038533 | *Brca1* | 0.74 | 0.033057 |
| *Ercc2* | 1.2 | 0.041554 | *Nol3* | 0.7 | 0.04037 |
| *Tnfrsf10b* | 1.63 | 0.045008 | *Egr1* | 0.55 | 0.040948 |
| *Ctgf* | 1.1 | 0.04758 | *Mgmt* | 0.83 | 0.043683 |
| *Eno1* | 1.36 | 0.049683 | *Xrcc3* | 0.86 | 0.044755 |
| | | | *Ppil2* | 0.88 | 0.045184 |
| | | | *Ppie* | 0.9 | 0.048768 |
| | | | *Tnfrsf11b* | 0.75 | 0.049312 |

### Table S1 – Differential Expression Gene Lists of targeted RNAseq

**Supplementary Table S1H – Differential Gene Expression of WT-CPZ vs APPPS1-CPZ**

| Symbol | Fold Change | adj. p-Value | Symbol | Fold Change | adj. p-Value |
|---|---|---|---|---|---|
| *Rb1* | 1.54 | 1.66061E-05 | *Ctsa* | 0.67 | 5.46188E-05 |
| *Xiap* | 1.37 | 6.31731E-05 | *Sirt2* | 0.83 | 0.000498423 |
| *Rap1a* | 1.45 | 8.13041E-05 | *Gadd45g* | 0.38 | 0.000581721 |
| *Sycp2* | 2.48 | 0.00013319 | *Xrcc1* | 0.8 | 0.00060194 |
| *Smug1* | 1.61 | 0.000170055 | *Mif* | 0.53 | 0.000949246 |
| *Rad18* | 1.43 | 0.000205468 | *Bbc3* | 0.71 | 0.001015096 |
| *Atr* | 1.57 | 0.000439004 | *Brca1* | 0.83 | 0.001210866 |
| *Rif1* | 1.64 | 0.000444179 | *Ddit4* | 0.53 | 0.001883707 |
| *Atm* | 1.45 | 0.000634496 | *Pgam2* | 0.69 | 0.002063495 |
| *Apaf1* | 2.18 | 0.000765516 | *Bmp1* | 0.46 | 0.00214972 |
| *Bmi1* | 1.45 | 0.000818183 | *Mms19* | 0.61 | 0.002290673 |
| *E2f1* | 1.32 | 0.001042813 | *Ppp1r15a* | 0.78 | 0.002389898 |
| *Ccnh* | 1.24 | 0.001489934 | *Aldoc* | 0.68 | 0.002555095 |
| *Mcl1* | 1.2 | 0.001610187 | *Pnkp* | 0.81 | 0.003106322 |
| *Ccng1* | 1.51 | 0.001628282 | *Cdc25b* | 0.64 | 0.003193596 |
| *Rev1* | 1.4 | 0.001748996 | *Ppil2* | 0.77 | 0.003194526 |
| *Cald1* | 1.38 | 0.001844269 | *Pim1* | 0.62 | 0.003843353 |
| *Tek* | 1.77 | 0.001848379 | *Xrcc3* | 0.7 | 0.003962897 |
| *Ccnc* | 1.47 | 0.001873173 | *Cpt2* | 0.68 | 0.004235657 |
| *Xrcc2* | 1.4 | 0.002043439 | *Rfx1* | 0.77 | 0.006626395 |
| *Sp1* | 1.2 | 0.002279162 | *Xab2* | 0.79 | 0.007234769 |
| *Terf1* | 1.56 | 0.002678689 | *Ccnf* | 0.78 | 0.007826727 |
| *Xrcc5* | 1.18 | 0.002696931 | *Sertad1* | 0.57 | 0.009781641 |
| *Vps13a* | 1.61 | 0.00274664 | *Mcm3* | 0.84 | 0.009867373 |
| *Mlh3* | 1.32 | 0.002823389 | *Msn* | 0.75 | 0.011275928 |
| *Pot1a* | 1.46 | 0.003156529 | *Rad9a* | 0.79 | 0.013107046 |
| *Tfdp2* | 1.21 | 0.003277162 | *Cd40lg* | 0.41 | 0.014010615 |
| *Mnat1* | 1.36 | 0.00334594 | *Mmp3* | 0.41 | 0.014010615 |
| *Rad21* | 1.24 | 0.003440183 | *Pgk2* | 0.41 | 0.014010615 |
| *Efnb2* | 1.28 | 0.004309619 | *Cib1* | 0.71 | 0.01533718 |
| *Timp1* | 1.52 | 0.004322032 | *Bax* | 0.63 | 0.015471245 |
| *Rad1* | 1.39 | 0.004490224 | *Mad2l2* | 0.88 | 0.017249263 |
| *Pdk3* | 1.1 | 0.004688622 | *Ercc1* | 0.78 | 0.017378926 |
| *Map3k2* | 1.24 | 0.00486861 | *Aldoa* | 0.71 | 0.017566656 |
| *Kpna2* | 2.02 | 0.00494824 | *Pdk2* | 0.71 | 0.019395251 |
| *Cdk5r1* | 1.32 | 0.005042964 | *G6pdx* | 0.88 | 0.021578402 |
| *E2f3* | 1.54 | 0.006084204 | *Hk3* | 0.72 | 0.023214134 |
| *Acsl4* | 1.44 | 0.00625717 | *Ruvbl2* | 0.69 | 0.02511066 |
| *Hgf* | 1.49 | 0.006950155 | *Lig1* | 0.73 | 0.025659011 |
| *Hk2* | 1.32 | 0.007154757 | *Nthl1* | 0.74 | 0.028858171 |
| *Cdh2* | 1.34 | 0.007395467 | *Efna1* | 0.56 | 0.03071962 |
| *Rad23a* | 1.14 | 0.007790167 | *Cdkn1a* | 0.78 | 0.030880135 |
| *Vegfc* | 1.47 | 0.009655269 | *Ccnd2* | 0.9 | 0.032117881 |
| *Cul1* | 1.21 | 0.010854905 | *Ldha* | 0.83 | 0.032729349 |
| *Runx2* | 2.16 | 0.012791422 | *Pfkfb3* | 0.87 | 0.034821279 |
| *Twist1* | 1.3 | 0.012813522 | *Tpi1* | 0.96 | 0.036760096 |
| *Pf4* | 1.47 | 0.013023351 | *Apex1* | 0.77 | 0.037267679 |
| *Bcl2* | 1.39 | 0.013121689 | *Vdac1* | 0.93 | 0.037426214 |
| *Skp2* | 1.26 | 0.01318376 | *Pkm* | 0.86 | 0.037751681 |
| *Hus1* | 1.1 | 0.013236611 | *Acvrl1* | 0.68 | 0.038698029 |
| *Rbbp8* | 1.19 | 0.015141994 | *Anapc2* | 0.88 | 0.039892529 |
| *Eno2* | 1.14 | 0.016301385 | *Thap3* | 0.81 | 0.040009659 |
| *Rad23b* | 1.32 | 0.016545879 | *Pcx* | 0.77 | 0.040324805 |
| *Wnt5a* | 1.75 | 0.017136738 | *Pfkm* | 0.85 | 0.04040667 |
| *Cdk7* | 1.36 | 0.018345971 | *Krt14* | 0.44 | 0.044215304 |
| *Ercc4* | 1.13 | 0.018867056 | *Gpd2* | 0.89 | 0.044665163 |
| *Cul2* | 1.21 | 0.019815127 | *Jmjd6* | 0.83 | 0.045024865 |
| *Tfrc* | 1.41 | 0.024920581 | *Ldhb* | 0.85 | 0.045636604 |
| *Eno1* | 1.42 | 0.026155662 | *Map2k3* | 0.8 | 0.045734852 |
| *Ets1* | 1.11 | 0.026531284 | *Rnf8* | 0.94 | 0.045999647 |
| *Nrp1* | 1.31 | 0.02672075 | *Ung* | 0.81 | 0.046884434 |
| *Nampt* | 1.37 | 0.028312078 | *Tep1* | 0.86 | 0.048997201 |
| *Cav2* | 1.28 | 0.029672763 | *Trp73* | 0.65 | 0.049052913 |
| *Nudt13* | 1.13 | 0.029750037 | *Ppie* | 0.88 | 0.050273788 |
| *Ocln* | 1.47 | 0.030912268 | *Xrcc6* | 0.87 | 0.050364795 |
| *Rad50* | 1.21 | 0.03192777 | | | |
| *Ercc6* | 1.18 | 0.033611852 | | | |
| *Mlh1* | 1.21 | 0.034878336 | | | |
| *Ptgs2* | 1.53 | 0.035524085 | | | |
| *Wee1* | 1.29 | 0.035568392 | | | |
| *Cul3* | 1.25 | 0.03865373 | | | |
| *Pfkfb1* | 1.35 | 0.039110246 | | | |
| *Rad17* | 1.16 | 0.039994646 | | | |
| *Mmp14* | 1.24 | 0.040357097 | | | |
| *Lig4* | 1.21 | 0.0417431 | | | |
| *Tgfbr1* | 1.31 | 0.044266361 | | | |
| *Kdr* | 1.49 | 0.045037522 | | | |
| *Atrx* | 1.21 | 0.047066061 | | | |
| *Ccnd1* | 1.37 | 0.048884092 | | | |

### Table S1 – Differential Expression Gene Lists of targeted RNAseq

#### Supplementary Table S1I – Differential Gene Expression of R140-CPZ vs APPPS1-CPZ

| Symbol | Fold Change | adj. p-Value | Symbol | Fold Change | adj. p-Value |
|---|---|---|---|---|---|
| *Rap1a* | 1.54 | 0.00018 | *Xab2* | 0.66 | 7.47446E-05 |
| *Mnat1* | 1.55 | 0.000266 | *Ercc1* | 0.83 | 0.000241892 |
| *Ccnc* | 1.43 | 0.000292 | *Gpi1* | 0.78 | 0.000262953 |
| *Smug1* | 1.45 | 0.000365 | *Msn* | 0.61 | 0.000295948 |
| *Terf1* | 1.64 | 0.000735 | *Aldoc* | 0.7 | 0.000377818 |
| *Rad23b* | 1.23 | 0.00096 | *Pkm* | 0.81 | 0.001540495 |
| *Rad50* | 1.31 | 0.001236 | *Map2k1* | 0.84 | 0.001664734 |
| *Uqcrfs1* | 1.21 | 0.00127 | *Cd40lg* | 0.3 | 0.001725468 |
| *Pf4* | 1.61 | 0.001635 | *Mmp3* | 0.3 | 0.001725468 |
| *Apaf1* | 1.68 | 0.002029 | *Pgk2* | 0.3 | 0.001725468 |
| *Rpa3* | 1.37 | 0.002238 | *Bmp1* | 0.42 | 0.001755882 |
| *Mlh3* | 1.39 | 0.00247 | *Vdac1* | 0.76 | 0.001838704 |
| *Bmi1* | 1.33 | 0.002788 | *Mad2l2* | 0.72 | 0.002039458 |
| *Atr* | 1.4 | 0.003093 | *Pcx* | 0.74 | 0.002050104 |
| *Birc2* | 1.32 | 0.003398 | *Anapc2* | 0.89 | 0.002285551 |
| *Cxcl12* | 1.55 | 0.003633 | *Pnkp* | 0.75 | 0.00231717 |
| *Tspan13* | 1.35 | 0.00368 | *Mms19* | 0.62 | 0.002418049 |
| *Mcl1* | 1.19 | 0.00409 | *Krt14* | 0.42 | 0.002663555 |
| *Sycp2* | 1.79 | 0.004201 | *Ccnt1* | 0.83 | 0.002930333 |
| *Desi1* | 1.15 | 0.004847 | *Sirt2* | 0.85 | 0.003181739 |
| *Cdc34* | 1.1 | 0.004939 | *Mcm5* | 0.78 | 0.003187084 |
| *Cdk8* | 1.2 | 0.005113 | *Sirt6* | 0.83 | 0.003529145 |
| *Mad2l1* | 1.29 | 0.005253 | *Xrcc1* | 0.79 | 0.004106321 |
| *Ccnh* | 1.18 | 0.005295 | *Pdk2* | 0.77 | 0.004516819 |
| *Rb1* | 1.3 | 0.006725 | *Mtor* | 0.85 | 0.004830446 |
| *Rad1* | 1.31 | 0.00757 | *Spata2* | 0.67 | 0.005074251 |
| *Pgk1* | 1.2 | 0.007753 | *Acly* | 0.79 | 0.005211429 |
| *Tek* | 1.45 | 0.008461 | *Adgrb1* | 0.82 | 0.005367316 |
| *Vegfc* | 1.77 | 0.009123 | *Ctsa* | 0.65 | 0.005400855 |
| *Bnip3* | 1.12 | 0.010068 | *Cdkn1b* | 0.87 | 0.005447824 |
| *Nampt* | 1.21 | 0.010234 | *Ldha* | 0.75 | 0.006209964 |
| *Polb* | 1.21 | 0.011699 | *Adm* | 0.72 | 0.008420799 |
| *Bccip* | 1.25 | 0.012675 | *Ercc2* | 0.81 | 0.008765065 |
| *Map3k2* | 1.19 | 0.01348 | *Ccnd3* | 0.8 | 0.008776765 |
| *Cul3* | 1.12 | 0.013572 | *Cdc20* | 0.64 | 0.010682897 |
| *Tmeff1* | 1.28 | 0.015773 | *Rtel1* | 0.47 | 0.010801889 |
| *Acsl4* | 1.29 | 0.017553 | *Edn1* | 0.5 | 0.010845918 |
| *Brca2* | 1.26 | 0.017891 | *Bcl2l1* | 0.83 | 0.010881496 |
| *Cul2* | 1.33 | 0.018334 | *Rnf8* | 0.87 | 0.012272556 |
| *Pinx1* | 1.36 | 0.019292 | *Ccnb1* | 0.8 | 0.0123841 |
| *Bnip3l* | 1.34 | 0.019362 | *Slc2a1* | 0.75 | 0.013218171 |
| *Ptges3* | 1.11 | 0.020097 | *Ccna2* | 0.76 | 0.013402421 |
| *Prkdc* | 1.46 | 0.02113 | *Ddit4* | 0.53 | 0.013639713 |
| *E2f3* | 1.13 | 0.022753 | *Cdc25b* | 0.59 | 0.013664259 |
| *Cdkn3* | 1.54 | 0.023057 | *Trp53* | 0.87 | 0.015219063 |
| *Xiap* | 1.16 | 0.024067 | *Aldoa* | 0.75 | 0.015652375 |
| *Vps13a* | 1.37 | 0.024126 | *Trp73* | 0.53 | 0.016205989 |
| *Tnfrsf11b* | 1.67 | 0.025285 | *Map2k3* | 0.75 | 0.016790254 |
| *Egr1* | 1.76 | 0.02696 | *Xrcc3* | 0.77 | 0.020719093 |
| *Cdh5* | 1.19 | 0.02837 | *Ruvbl2* | 0.63 | 0.020915249 |
| *Aurkb* | 2.19 | 0.028897 | *Cib1* | 0.83 | 0.020984933 |
| *Ccnd1* | 1.32 | 0.030099 | *Tbx2* | 0.65 | 0.021325762 |
| *Fen1* | 1.11 | 0.030637 | *Cpt2* | 0.77 | 0.021333354 |
| *Ccng1* | 1.34 | 0.031335 | *Ppp1r15a* | 0.79 | 0.021652886 |
| *Fam162a* | 1.16 | 0.033184 | *Nthl1* | 0.62 | 0.022849754 |
| *Bcl2* | 1.45 | 0.033514 | *Cdk2* | 0.73 | 0.023362175 |
| *Xrcc2* | 1.4 | 0.036066 | *Pim1* | 0.56 | 0.025013891 |
| *Cul1* | 1.14 | 0.036598 | *Mif* | 0.7 | 0.025767302 |
| *Cald1* | 1.13 | 0.036839 | *Atp5a1* | 0.85 | 0.025983712 |
| *Snai2* | 1.27 | 0.03812 | *Angptl4* | 0.38 | 0.027460075 |
| *Rad17* | 1.15 | 0.038454 | *Adora2b* | 0.85 | 0.028698635 |
| *Lig4* | 1.24 | 0.042766 | *Rpa1* | 0.88 | 0.031906413 |
| *Ptgs2* | 1.55 | 0.043708 | *Pdk1* | 0.9 | 0.033549264 |
| *Atm* | 1.16 | 0.044089 | *Hk1* | 0.81 | 0.035699613 |
| *Nop10* | 1.26 | 0.044694 | *Bax* | 0.75 | 0.035728165 |
| *Kpna2* | 1.18 | 0.045583 | *Rfx1* | 0.76 | 0.036019506 |
| *Dll4* | 1.37 | 0.047876 | *Tnks* | 0.82 | 0.037393962 |
| *Col1a2* | 1.19 | 0.048745 | *Slc2a3* | 0.76 | 0.038850346 |
| | | | *Casp6* | 0.86 | 0.039242101 |
| | | | *Xpc* | 0.9 | 0.040293911 |
| | | | *Tinf2* | 0.85 | 0.040639982 |
| | | | *Ppil2* | 0.83 | 0.044279856 |
| | | | *Txnip* | 0.6 | 0.045298986 |
| | | | *Eno2* | 0.88 | 0.046824966 |
| | | | *Lig1* | 0.72 | 0.047379344 |
| | | | *Gadd45g* | 0.58 | 0.047720615 |
| | | | *Odc1* | 0.88 | 0.048554959 |
| | | | *Bbc3* | 0.74 | 0.048556645 |
| | | | *Zfp446* | 0.71 | 0.048977571 |
| | | | *Ets2* | 0.84 | 0.049660642 |

### Table S1 – Differential Expression Gene Lists of targeted RNAseq

**Supplementary Table S1J – Gene list of QIAseq Targeted RNA Mouse Cancer Transcriptome Panel (Qiagen RMM-003Z)**

| Apoptosis | Cell Cycle | Cell Senescence | DNA Damage & Repair | Epithelial-to-Mesenchymal Transition | Hypoxia | Metabolism | Telomeres & Telomerase | Angiogenesis |
|---|---|---|---|---|---|---|---|---|
| Abl1 | Abl1 | Atm | Apex1 | Ahnak | Adm | Acly | Acd | Acvrl1 (Alk1) |
| Akt1 | Anapc2 | Bmi1 | Atrx | Bmp1 | Adora2b | Acsl4 | Dkc1 | Adgrb1 |
| Apaf1 | Atm | Ccnd1 | Bbc3 | Cald1 | Aldoc | Aldoa | Gar1 | Akt1 |
| Atp6v1g2 | Atr | Ccne1 | Blm | Camk2n1 | Angptl4 | Aldob | Hsp90aa1 | Ang |
| Bax | Aurka | Cdk2 | Brip1 | Cav2 | Ankrd37 | Atp5a1 | Mre11a | Angpt1 |
| Bcl2 | Aurkb | Cdk4 | Cib1 | Cdh1 (E-cadherin) | Anxa2 | Cox5a | Nbn (Nbs1) | Angpt2 |
| Bcl2a1a (Bfl-1 A1) | Bccip | Cdk6 | Ddb1 | Cdh2 (N-cadherin) | Arnt | Cpt2 | Nhp2 | Ccl2 (MCP-1) |
| Bcl2a1b | Bcl2 | Cdkn1a (p21Cip1 Waf1) | Ddb2 | Col1a2 | Atr | Eno1 | Nop10 | Ccn2 |
| Bcl2a1d | Birc5 | Cdkn2a (p16INK4a) | Ercc1 | Col3a1 | Bhlhe40 (Dec1) | Eno2 | Obfc1 | Cdh5 |
| Bcl2l1 (Bcl-XL) | Brca1 | Cdkn2d (p19INK4d) | Ercc2 (Xpd) | Col5a2 | Blm | Fbp1 | Pinx1 | Col18a1 |
| Bcl2l11 | Brca2 | Chek1 | Ercc3 (Xpb) | Desi1 | Bnip3 | Fbp2 | Pot1a | Col4a3 |
| Birc2 (cIAP1, cIAP2) | Casp3 | Chek2 (Rad53) | Ercc4 | Dsp | Bnip3l | G6pc | Ptges3 | Cxcl12 |
| Birc3 (cIAP1, cIAP2) | Ccna2 | E2f1 | Ercc5 | Fgfbp1 | Car9 | G6pdx | Rad50 | Dll4 |
| Casp1 (Ice) | Ccnb1 | E2f3 | Ercc6 | Fn1 | Ccng2 | Gpd2 | Rap1a | Edn1 |
| Casp2 | Ccnb2 | Ets1 | Ercc8 | Foxc2 | Ctsa | Gpi1 | Rif1 | Efna1 |
| Casp3 | Ccnc | Ets2 | Exo1 | Gng11 | Ddit4 (Redd1) | Hk1 | Rtel1 | Efnb2 |
| Casp6 | Ccnd1 | Igfbp5 | Fen1 | Gsc | Dnajc5 | Hk2 | Sirt2 | Egf |
| Casp7 | Ccnd2 | Igfbp7 | Gadd45g | Igfbp4 | Edn1 | Hk3 | Sp1 | Eng (Evi-1) |
| Casp9 | Ccnd3 | Ing1 | Lig1 | Il1rn | Egr1 | Ldha | Tep1 | Ephb4 |
| Cd40 | Ccne1 | Map2k1 (Mek1) | Lig3 | Itga5 | Eif4ebp1 | Ldhb | Terf1 | F3 |
| Cd40lg | Ccnf | Map2k3 (Mek3) | Lig4 | Itgav | Epo | Lpl | Terf2 | Fgf1 |
| Cflar (Casper) | Ccng1 | Mdm2 | Mgmt (Agt) | Krt14 | F3 | Mtor | Terf2ip | Fgf2 (bFGF) |
| Cyld | Ccng2 | Rb1 | Mlh1 | Krt19 | Fam162a | Pck1 | Tert | Flt1 (Vegfr1) |
| Dffa | Ccnh | Rbl2 | Mlh3 | Mmp2 | Gpi1 | Pck2 | Tinf2 | Fn1 |
| Fas | Ccnt1 | Serpinb2 (Pai-2) | Mms19 | Mmp3 | Hk2 | Pcx | Tnks (Tin1) | Hgf |
| Fasl | Cdc16 | Sirt1 | Mre11a | Mmp9 | Hmox1 | Pdha1 | Tnks2 | Hif1a |
| Gadd45a | Cdc20 | Tbx2 | Msh2 | Msn | Ier3 | Pdk1 | | Id1 |
| Igf1r | Cdc25a | Trp53 (p53) | Msh3 | Mst1r (Ron) | Igfbp3 | Pdk2 | | Ifng |
| Mcl1 | Cdc25b | Twist1 | Msh6 | Nudt13 | Jmjd6 | Pdk3 | | Igf1 |
| Nol3 | Cdc25c | | Neil1 | Ocln | Ldha | Pdk4 | | Il1b |
| Spata2 | Cdc34 | | Neil3 | Rgs2 | Lgals3 | Pfkfb1 | | Il6 |
| Sycp2 | Cdc6 | | Nthl1 | Serpine1 (Pai-1) | Lox | Pfkfb2 | | Itgb3 |
| Tnf | Cdk1 (Cdc2) | | Parp1 | Snai1 (Snail) | Map3k1 (Mekk1) | Pfkfb3 | | Jag1 |
| Tnfrsf10b (Dr5) | Cdk2 | | Parp2 | Snai2 | Met | Pfkfb4 | | Kdr (Vegfr2) |
| Tnfrsf11b (Opg) | Cdk4 | | Parp3 | Snai3 | Mif | Pfkl | | Lep (Leptin) |
| Tnfrsf1a (Tnfr1) | Cdk5r1 | | Pms1 | Sox10 | Mmp9 | Pfkm | | Mmp14 |
| Traf2 | Cdk5rap1 | | Pms2 | Sparc | Mxi1 | Pgam2 | | Mmp2 |
| Trp53 (p53) | Cdk6 | | Pnkp | Spp1 | Nampt | Pgk1 | | Mmp9 |
| Xiap | Cdk7 | | Polb | Steap1 | Ndrg1 | Pgk2 | | Nos3 (eNOS) |
| | Cdk8 | | Poll | Tcf4 | Nos3 (eNOS) | Pgm1 | | Notch4 |
| | Cdkn1a (p21Cip1 Waf1) | Ppp1r15a (Gadd34) | Tfpi2 | Odc1 | Pklr | | Nrp1 | |
| | Cdkn1b (p27Kip1) | Prkdc | Timp1 | P4ha1 | Pkm | | Nrp2 | |
| | Cdkn2a (p16INK4a) | Rad18 | Tmeff1 | Pfkfb4 | Prkaa1 (Ampk) | Pf4 | | |
| | Cdkn3 | | Rad23a | Tspan13 | Pgk1 | Runx2 | | Pgf |
| | Chek1 | | Rad23b | Twist1 | Pim1 | Sirt6 | | Plg |
| | Chek2 (Rad53) | Rad50 | Vcan | Plau (uPA) | Slc2a1 | | Ptgs2 (Cox2) | |
| | Cks1b | | Rad51 | Vim | Ruvbl2 | Slc2a4 (Glut4) | Serpine1 (Pai-1) | |
| | Cks2 | | Rad51b | Vps13a | Serpine1 (Pai-1) | Stk11 (Lkb1) | | Serpinf1 |
| | Cul1 | | Rad52 | Wnt5a | Slc16a3 | Tpi1 | | Tek (Tie-2) |
| | Cul2 | | Rad54l | Wnt5b | Slc2a1 | Uqcrfs1 | | Tgfb1 |
| | Cul3 | | Rev1 | | Slc2a3 | | | Tgfbr1 (ALK5) |
| | E2f1 | | Rnf8 | | Tfrc | | | Thbs1 (TSP-1) |
| | E2f4 | | Rpa1 | | Txnip | | | Thbs2 |
| | Gadd45a | | Rpa3 | | Vdac1 | | | Timp2 |
| | Gtse1 | | Smug1 | | Vegfa | | | Tnf |
| | Hus1 | | Trp53bp1 | | | | | Vegfa |
| | Kntc1 | | Trp73 | | | | | Vegfc |
| | Kpna2 | | Ung | | | | | Vegfd (Vegfd) |
| | Mad2l1 | | Xab2 | | | | | |
| | Mad2l2 | | Xpa | | | | | |
| | Mcm2 | | Xpc | | | | | |
| | Mcm3 | | Xrcc1 | | | | | |
| | Mcm4 | | Xrcc2 | | | | | |
| | Mcm5 | | Xrcc3 | | | | | |
| | Mdm2 | | Xrcc4 | | | | | |
| | Mki67 | | Xrcc5 | | | | | |
| | Nbn (Nbs1) | | | | | | | |
| | Plk1 | | | | | | | |
| | Rad1 | | | | | | | |
| | Rad17 | | | | | | | |
| | Rad9a | | | | | | | |
| | Rb1 | | | | | | | |
| | Rbbp8 | | | | | | | |
| | Rbl1 | | | | | | | |
| | Rbl2 | | | | | | | |
| | Sertad1 | | | | | | | |
| | Skp2 | | | | | | | |
| | Stmn1 | | | | | | | |
| | Tfdp1 | | | | | | | |
| | Tfdp2 | | | | | | | |
| | Trp53 (p53) | | | | | | | |
| | Wee1 | | | | | | | |

**Table S1. Differential Gene Expression Lists of targeted RNAseq**

DEGs from all pairwise comparison are listed in table S1A to S1I with blue list indicates significant upregulation and red list indicates significant downregulation across all samples. Fold changes and adjusted P values are listed in each DEG. The full gene list of QIAseq Targeted RNA Mouse Cancer Transcriptome Panel is listed in table S1J.
