## Supplementary material for "Cuprizone drives divergent neuropathological changes in different mouse models of Alzheimer’s disease": Fig. S2

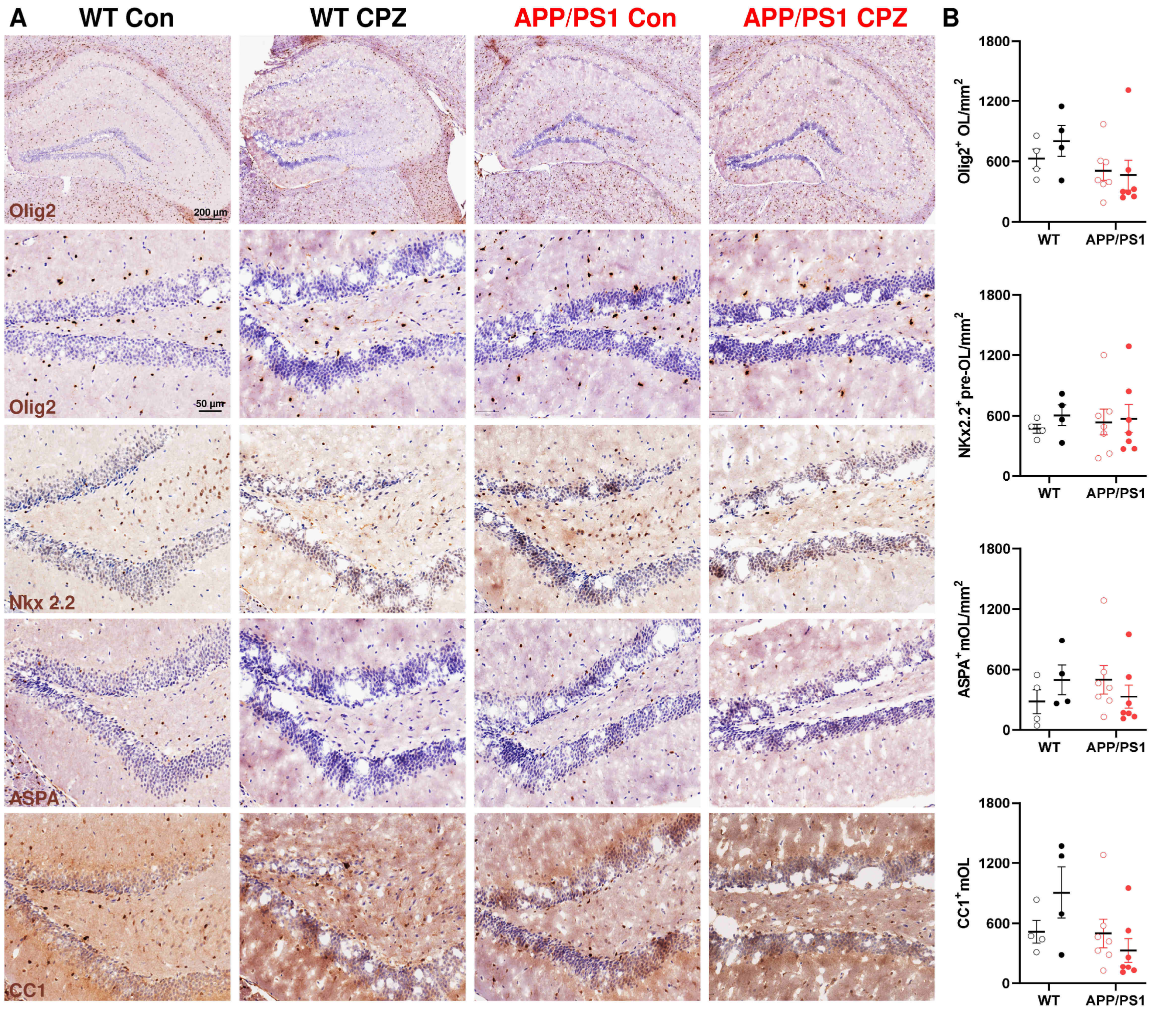


**Figure S2. No significant oligodendrocyte pathology in the hippocampus**

**A** Representative image of Olig2 the hippocampus at Bregma -1.3 (*upper*), and representative image of Olig2, Nkx 2.2, ASPA and CC1 for all groups at middle magnifications (lower). **B** No significant differences in Olig2$^{+}$, Nkx 2.2$^{+}$, ASPA$^{+}$ and CC1$^{+}$ OLs between genotype nor treatment were found in the hippocampus.
