## Supplementary material for "Cuprizone drives divergent neuropathological changes in different mouse models of Alzheimer’s disease": Fig. S1

### SUPPLEMENTARY DATA

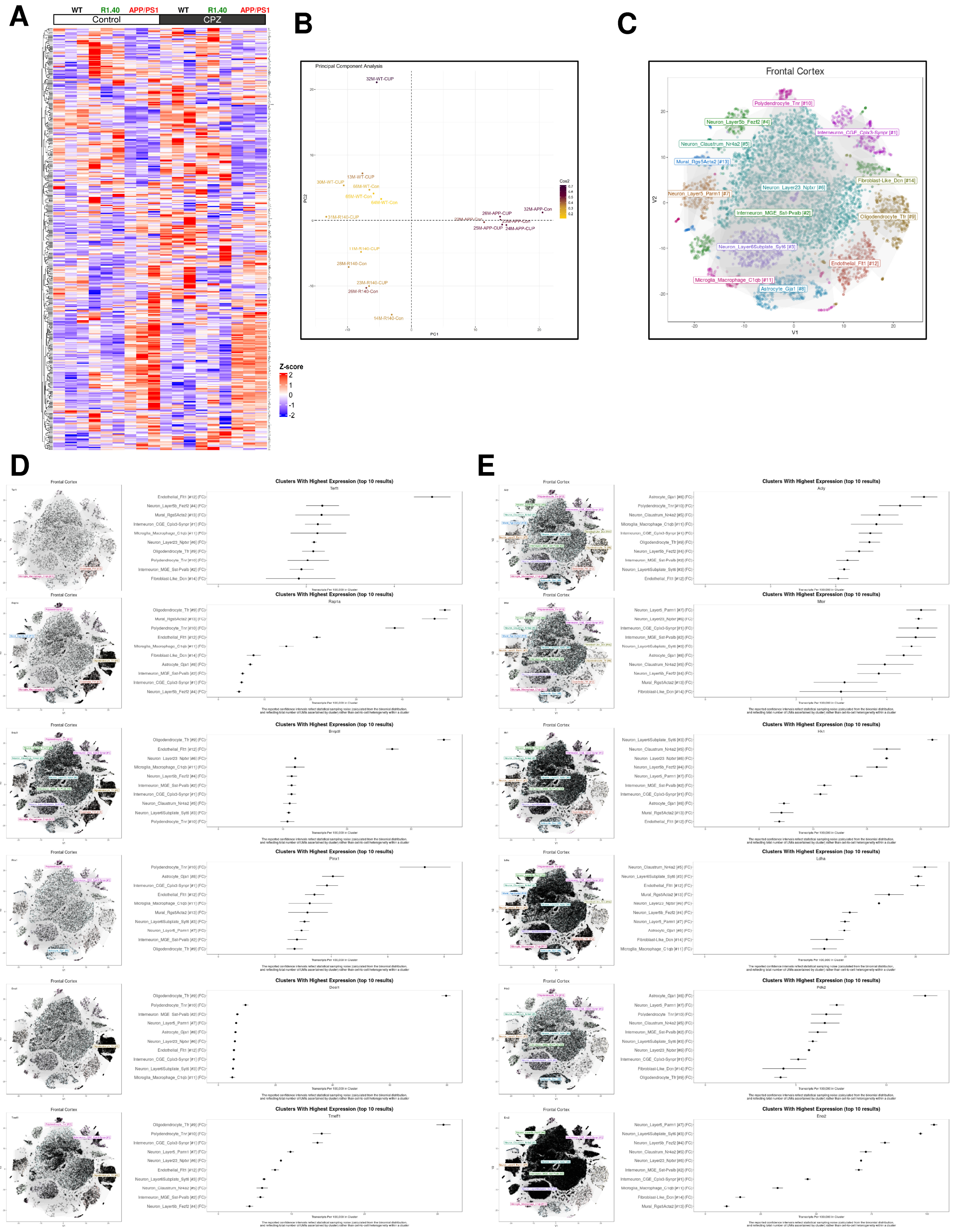


**Figure S1. Extended data for targeted RNAseq of cerebral cortex**

**A** Heatmap of the normalized expression of all genes in the QIAseq Targeted RNA Mouse Cancer Transcriptome Panel (Qiagen, Germany, RMM-003Z) among WT, R1.40 and APP/PS1 with or without cuprizone treatment (Control, CPZ). **B** Principal component analysis showing the segregation of samples by genotype and cuprizone treatment. **C** Frontal cortex tSNE map from a mouse brain single cell RNA-seq database (Dropviz (72) ) with the indications for each cell subtypes. The cellular expression of transcripts in frontal cortex derived from **D** *Establishment of localization in cell*, (*Terf1*, *Rap1a*, *Bnip3l*, *Pinx1*, *Desi1*, *Tmeff1*, GO:0051649) and, **E** *Small molecule metabolic process*, (*Acly*, *Mtor*, *Hk1*, *Ldha*, *Pdk2*, *Eno2*, GO:0044281) are shown. The majority in the former, but not the latter, is enriched in oligodendrocyte lineage.
